# Structure of Dimeric Lipoprotein Lipase Reveals a Pore for Hydrolysis of Acyl Chains

**DOI:** 10.1101/2023.03.21.533650

**Authors:** Kathryn H. Gunn, Saskia B. Neher

**Author notes:** Corresponding author: Saskia B. Neher. Author Contributions: K.H.G. and S.B.N. planned the experiments and wrote the manuscript. K.H.G. performed experiments and analyzed data. Competing Interest Statement: The authors declare no competing interests.

## Abstract

Lipoprotein lipase (LPL) hydrolyzes triglycerides from circulating lipoproteins, releasing free fatty acids. Active LPL is needed to prevent hypertriglyceridemia, which is a risk factor for cardiovascular disease (CVD). Using cryogenic electron microscopy (cryoEM), we determined the structure of an active LPL dimer at 3.9 Å resolution. This is the first structure of a mammalian lipase with an open, hydrophobic pore adjacent to the active site. We demonstrate that the pore can accommodate an acyl chain from a triglyceride. Previously, it was thought that an open lipase conformation was defined by a displaced lid peptide, exposing the hydrophobic pocket surrounding the active site. With these previous models after the lid opened, the substrate would enter the active site, be hydrolyzed and then released in a bidirectional manner. It was assumed that the hydrophobic pocket provided the only ligand selectivity. Based on our structure, we propose a new model for lipid hydrolysis, in which the free fatty acid product travels unidirectionally through the active site pore, entering and exiting opposite sides of the protein. By this new model, the hydrophobic pore provides additional substrate specificity and provides insight into how LPL mutations in the active site pore may negatively impact LPL activity, leading to chylomicronemia. Structural similarity of LPL to other human lipases suggests that this unidirectional mechanism could be conserved but has not been observed due to the difficulty of studying lipase structure in the presence of an activating substrate. We hypothesize that the air/water interface formed during creation of samples for cryoEM triggered interfacial activation, allowing us to capture, for the first time, a fully open state of a mammalian lipase. Our new structure also revises previous models on how LPL dimerizes, revealing an unexpected C-terminal to C-terminal interface. The elucidation of a dimeric LPL structure highlights the oligomeric diversity of LPL, as now LPL homodimer, heterodimer, and helical filament structures have been elucidated. This diversity of oligomerization may provide a form of regulation as LPL travels from secretory vesicles in the cell, to the capillary, and eventually to the liver for lipoprotein remnant uptake. We hypothesize that LPL dimerizes in this active C-terminal to C-terminal conformation when associated with mobile lipoproteins in the capillary.

## Introduction

Lipases are of critical importance due to their roles liberating free fatty acids (FFA), enabling their transport across membranes. Lipoprotein lipase (LPL) is a secreted enzyme that is active in the capillaries, where it hydrolyzes triglycerides found in chylomicrons and very low-density lipoproteins (VLDL) to release FFA, which are transported across the capillary endothelial membrane as a source of energy (1). In addition to acting on triglycerides, LPL has phospholipase activity (2), making it essential to regulate where and when LPL is active to prevent off-target hydrolysis events. LPL is synthesized in the adipose and oxidative tissues and is packaged into secretory vesicles using either a sphingomyelin specific pathway with heparan sulfate proteoglycan (HSPG) syndecan-1 (SDC1) or a bulk flow secretion pathway (3). LPL secretion is regulated in part by nutritional and metabolic cues from the environment, and some LPL is stored in vesicles until nutritional signaling induces secretion (4). We have previously shown that in the storage vesicles, LPL adopts a filament-like structure that follows the vesicle membrane and resembles the helical oligomer of LPL we solved by cryoEM (5). Following exocytosis, LPL is released from the SDC1 HSPGs and transits the interstitial space to reach GPIHBP1, which is expressed by capillary endothelial cells (6). GPIHBP1 is a glycolipid anchored membrane protein with an extracellular binding domain for LPL. LPL forms a 1:1 complex with GPIHBP1 (7, 8). GPIHBP1 transports LPL into the capillary, where it can interact with triglyceride rich lipoproteins (TRLs) to release FFAs (9). LPL can also be found free from GPIHBP1 and associated with mobile TRLs in the blood (10). Independent from its role hydrolyzing triglycerides, LPL on TRLs mediates lipoprotein remnant uptake (11, 12).

Historically, LPL was thought to have two oligomeric states: an active dimer and inactive monomer (13, 14). Recent structural work has challenged this paradigm. First, two crystal structures of LPL in complex with GPIHBP1 were solved (7, 8). The LPL/GPIHBP1 heterodimer formed a heterotetramer in the asymmetric unit of the crystal, but this conformation occluded access to the active site and further analysis suggested it was likely a crystallization artefact (7). Size exclusion multi-angle light scattering (SEC-MALS) was used to show that in solution LPL/GPIHBP1 exists as a 1:1 heterodimer (7). This work confirmed that the LPL/GPIHBP1 heterodimer is an active form of LPL. It has also been observed that monomers of LPL may be active in the absence of GPIHBP1 (15). Next, an inactive helical oligomer of LPL was identified by cryoEM, showing that there were also other inactive forms of LPL beyond a monomer (5). These studies left the structure of the LPL dimer unresolved. Studies using radiation inactivation, ultracentrifugation, ELISA sandwich assays, and single molecule fluorescence resonance energy transfer (smFRET) indicate the existence of an LPL dimer (13, 14, 16–18). A shared feature of these experiments that showed the existence of dimeric LPL was that they were performed without GPIHBP1.

Structures of many mammalian lipases have been solved, including pancreatic triacylglycerol lipase (19–21), pancreatic lipase related protein 1 (22) and 2 (23), monoacylglycerol lipase (24), lipoprotein lipase (5, 7, 8), gastric lipase (25), bile-salt activated lipase (26, 27), and lysosomal acid lipase (28). A few of these structures were solved in open conformations where the lid peptide residues were shifted to expose a hydrophobic pocket adjacent to the active site (20, 21, 23). Lipases achieve an open state by undergoing interfacial activation, which occurs when the lipase associates with a nonpolar-aqueous interface (29). The combination of the lid peptide and hydrophobic pocket is thought to provide mammalian lipases substrate selectivity. An outlier in the eukaryotic lipase family is found in fungi, specifically Candida rugosa lipase (30), which uses a tunnel for substrate binding rather than a hydrophobic pocket. Structures have shown acyl chain analogs are able to enter and occupy this hydrophobic tunnel (30). The tunnel controls entrance to the active site and a further pocket beyond it. Subsequent mutational studies have shown that mutations made in this tunnel can alter substrate specificity (31).

We set out to solve the structure of LPL in the absence of GPIHBP1, hypothesizing that this would reveal an active LPL dimer. Indeed, we were able to solve the structure of a dimeric LPL oligomer, which revealed a series of novel insights, including those with mechanistic implications for LPL and mammalian lipases in general.

## Results

### LPL treated with deoxycholate prevents formation of inactive helices

To generate a homogenous sample of active LPL for cryoEM, we needed to disrupt the inactive LPL helices that form at high protein concentrations. We previously found that treatment with the bile salt deoxycholate causes dissolution of LPL helices (5), and LPL treated with deoxycholate is stable and active (32). Therefore, we treated LPL with deoxycholate and dialyzed it into buffer both with and without additional deoxycholate. Deoxycholate has a critical micelle concentration (CMC) that can vary based on temperature, but the concentration we used (1 mM) is below the CMC and can be removed by dialysis (33). We analyzed both conditions by negative stain transmission electron microscopy (nsTEM) and cryoEM and found LPL helices were not present in either. These data show that binding to deoxycholate and helix dissolution is not affected by subsequent dialysis to a buffer without deoxycholate. We also tested the activity of LPL by monitoring the release of FFA from VLDL using an enzyme coupled fluorescent assay (34). Interestingly, LPL activity was slightly diminished when deoxycholate was present compared to LPL alone. We found that the activity of deoxycholate-treated LPL subsequently dialyzed with or without deoxycholate was comparable (Supplemental Figure 1A&B). Bovine LPL, which shares ∼94% sequence identity with human LPL (35), was used for the experiments described in this work unless otherwise specified.

### LPL forms a C-terminal to C-terminal homodimer

After confirming we had a homogenous, active preparation of LPL (Supplemental Figure 1C), we analyzed the sample using single particle cryoEM (Supplemental Figure 2A). Initial 2D classification revealed one clear class of LPL with secondary structure visible (Figure 1A). However, few other classes were present in the data. This result suggested that LPL was adopting a preferred orientation by interacting with the air/water interface (AWI) during grid freezing (36, 37). Multiple methods were attempted to alleviate the preferred orientation via detergents and additives. None of these were able to disrupt the association of LPL with the AWI. We ultimately were able to overcome the preferred orientation despite the relatively small size of our protein (∼100 kDa dimer) by using a tilting strategy (Figure 1A) combined with specific enrichment of rare particle views using the Topaz neural network (38) (Supplemental Figure 2B). We collected data sets at 3 tilts: 0°, 30°, and 45°. We processed each tilted data set individually using cryoSPARC (39) and focused on enriching for rare 2D classes by iteratively training Topaz (38) to pick non-preferred orientation particles for each tilt set. We then combined the particles from all tilts.

**Figure 1.**
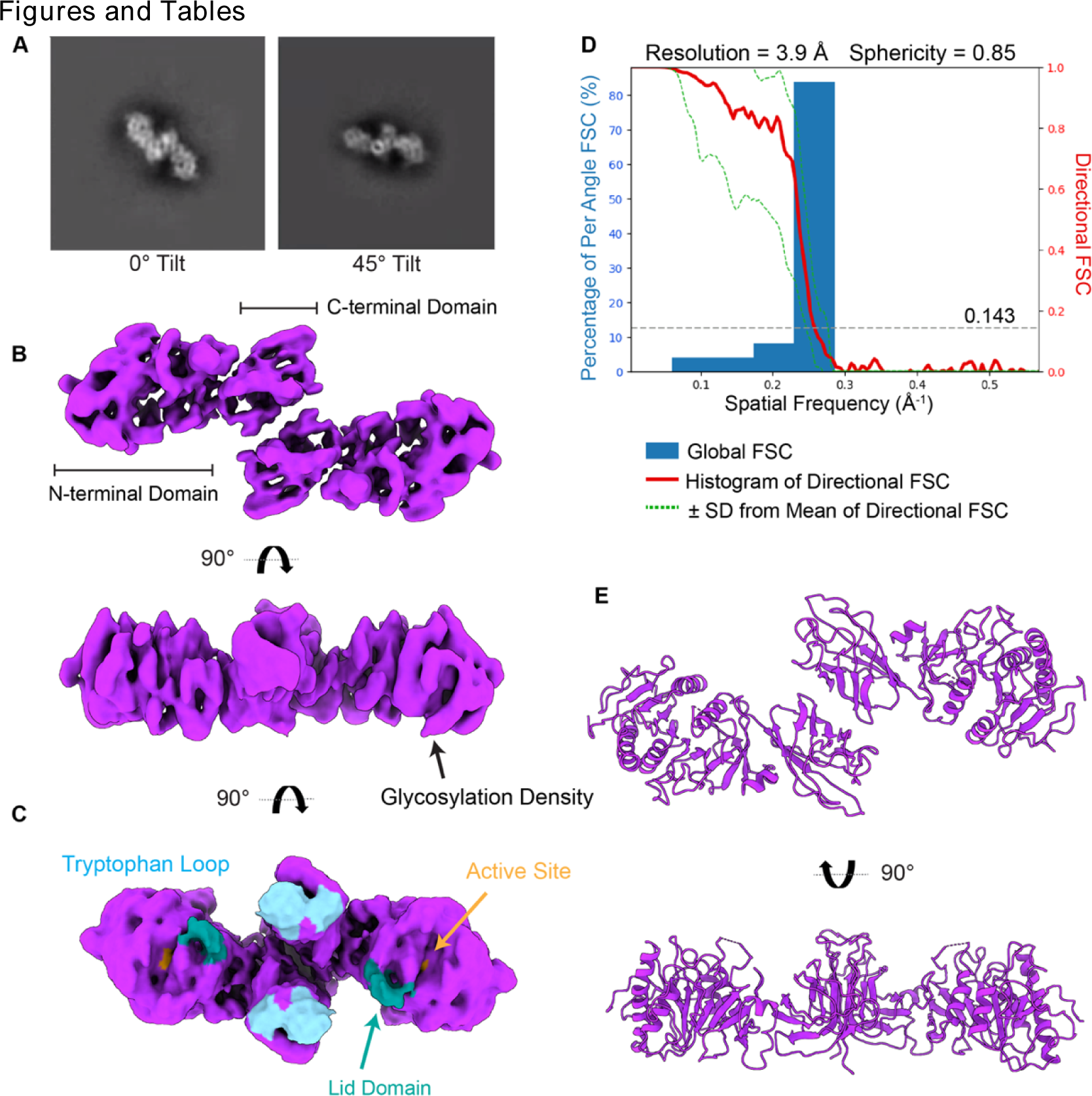
LPL forms C-terminal to C-terminal homodimers. A) The 2D class of LPL’s preferred orientation at 0° (left). Tilting the grid to 45° allowed us to capture an alternate view of this orientation (right). B) The density map for the LPL dimer. The dimerization interface is at the C-terminus of both LPL subunits. Density corresponding to the glycosylation of N73 was observed. C) When thresholding of the density map is increased, density for the lid domain of LPL appears (aqua). The tryptophan loop domain in the C-terminus, which is involved in substrate recognition, can be seen along the same plane as the density corresponding to the lid peptide (cyan). The lid peptide is likely too mobile to be resolved at high resolution. The lid domain density has shifted downward and to the side of the active site (orange) adopting an open conformation. D) The resolution of the LPL dimer is 3.9 Å. The density map was analyzed using the 3DFSC server revealing a sphericity of 0.85, which indicates that we were able to create a nearly complete reconstruction, despite anisotropy resulting from the preferred orientation of the particles (Supplemental Figure 3B). E) The molecular map of the LPL dimer was built into the density map (Supplemental Figure 4E). The residues of the lid domain were not modeled because their location was not determined with high resolution.

We found that the LPL dimer had C2 symmetry, which was applied after initial rounds of C1 symmetry refinement (Figure 1B). The final resolution was 3.9 Å using the gold-standard Fourier shell correlation (GSFSC) with a cut-off of 0.143 on the directional FSC (Figure 1D). Analysis with the 3DFSC server (36) indicated that although anisotropy was still present in the structure, the sphericity was 0.85 out of 1, highlighting the success of a combined tilting and Topaz picking. The distribution of particles from each view of the final structure is illustrated in Supplemental Figure 3A. We were able to unambiguously dock LPL into the density, identify secondary structure, observe separation of β-sheets (Supplemental Figure 2C), and see density for large amino acid side chains. A map of LPL was built into the density using Coot and Phenix (Figure 1E, Table 1). LPL has a flexible lid peptide that covers its active site, and we were unable to build the map to fit the LPL lid density at 3.9 Å, although it can be clearly seen when the thresholding is reduced (Figure 1C). The correlation between the map and model is 0.77 (Table 1). Due to the residual anisotropy of the model, there is lower confidence for the fit of some amino acid side chains, leading to the lower correlation value. The interior residues of the protein were at higher resolution than the outer residues (Supplemental Figure 3B).

**Table 1.** CryoEM Validation Statistics.

| **Data Collection and Processing** |  |
| --- | --- |
| Magnification | 45,000 |
| Voltage (kV) | 200 |
| Pixel size (Å) | 0.88 |
| Total Dose (e<sup>-</sup>/Å<sup>2</sup>) | 55.1 |
| Number of Particles | 527,205 |
| B-factor | -100 |
| **Map Resolution (Å)** |  |
| Model:map FCR (0.5) | 4.5 |
| Map:map FSC (0.143) | 3.9 |
| d<sub>99</sub> | 4.6 |
| Symmetry | C2 |
| **Model Refinement** |  |
| Model to map CC | 0.77 |
| Clash score, all atoms | 4.84 |
| Ramachandran favored (%) | 90.51 |
| Ramachandran outliers (%) | 0 |
| Rotamer outliers (%) | 0.27 |
| C<sub>β</sub> deviations > 0.25 Å | 0 |
| **RMS Deviations** |  |
| Bond (Å) | 0.005 |
| Angles (°) | 1.277 |
| **Deposition IDs** |  |
| PDB ID | 8ERL |
| EMDB ID | 28554 |

Unexpectedly, the structure shows that LPL forms a homodimer associated at the C-terminal domains (“tails”) of both subunits (Figure 1). Intriguingly, in the tail-to-tail dimer, two features of LPL which are key for substrate interaction are situated along the same surface. Specifically, the C-terminal tryptophan-loop, which helps recognize TRL substrates (40), and the N-terminal lid peptide, which opens in response to a lipid-water interface, are arrayed on a single plane (Figure 1C).

### Active site pore in the LPL dimer structure

The active LPL dimer is oriented with both its substrate recognizing tryptophan loops and active sites facing the same direction, suggesting an active conformation that could engaged with a TRL substrate (Figure 2A). We thus analyzed the LPL active site in the structure and found a clearly defined pore traversing LPL which is directly adjacent to the active site residues. The fit of the pore lining residues to the cryoEM map can be seen in Supplemental Figure 4. This open pore in LPL is facilitated by shifts of many amino acids, these differences are illustrated by morphing other LPL structures to the LPL from our homodimer (Supplemental Movie 1-3).

**Figure 2.**
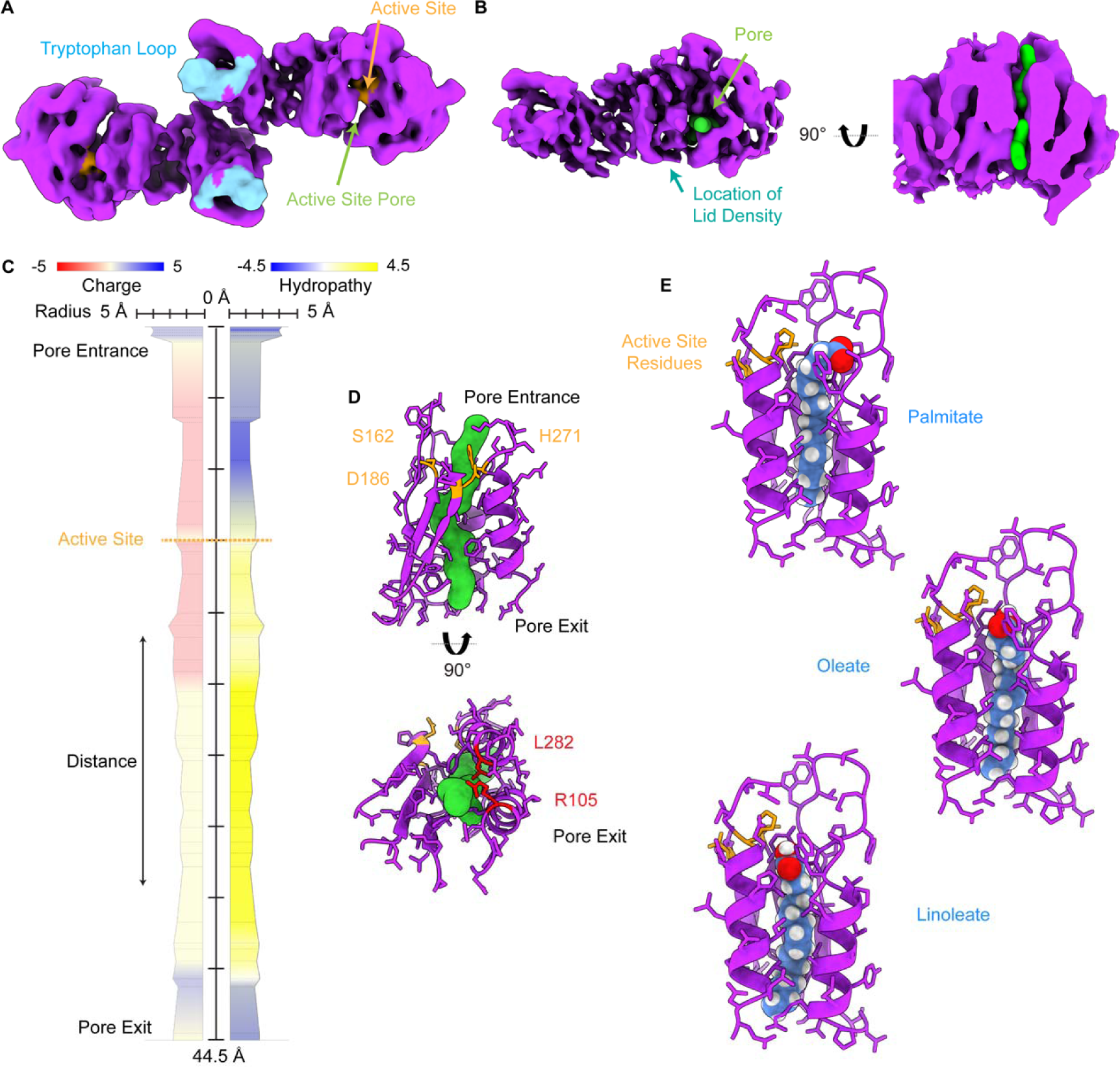
The LPL homodimer reveals an open active site pore. A) The LPL dimer map colored to show the LPL active site residues (orange) and the tryptophan loop (cyan), which is involved in substrate recognition. An area without protein density is seen in the N-terminal of LPL suggested the existence of a pore spanning the protein. B) We analyzed the LPL model and were able to identify a pore that spanned the N-terminal domain (green), shown here in the density map. The pore is located adjacent to the active site and an aqua arrow indicates the position of the open lid density, which has swung to the side of this pore. A cutaway of the density shows that the pore spans the N-terminal domain of LPL. C) MOLEonline was used to create a trace of the characteristics of the amino acids that line the identified pore. The residues corresponding to the entrance to the LPL pore are at the top of the diagram and the path of the pore travels to the terminus at the bottom. The pore is 44.5 Å in length with a bottleneck radius of 1.4 Å. D) The residues surrounding the pore are displayed to show how the pore interacts with surrounding amino acids. The active site residues (S162, D186, and H271) are indicated in orange. H271 directly abuts the pore density. Looking at the pore terminus (the end opposite of the LPL lid domain), we can see that there is an unobstructed exit from the pore. Two residues, L282 and R105, known to cause LPL deficiency in humans are located near the pore terminus (red). Models of these mutations can be seen in Supplemental Figure 9. E) A sphere representation of 3 common acyl chains fit into the hydrophobic pore was created with PyRosetta. Ligands shown are saturated fatty acid palmitate (C16:0), monounsaturated oleate (C18:1), and doubly unsaturated lineolate (C18:2). The starting and final models for these fits are available as supplemental materials, including the results of the PyRosetta refinement in the pdb files.

We used MOLEonline (41), an automated toolkit for analysis of tunnels and pores, to identify the active site pore using our protein model. A pore with a length of ∼45 Å was identified that passes through the N-terminal domain of LPL (Figure 2B-D). We also performed this analysis on LPL from the helical LPL oligomer, LPL/GPIHBP1, and other mammalian lipase structures. No comparable pores were identified. The pore in the dimer structure has a bottleneck with a radius of 1.4 Å, which is sufficient for the carbon hydrogen backbone of a fatty acid. The pore has a highly hydrophobic section that could accommodate a fatty acid tail, potentially aligning the hydrolysable ester bond with the active site residues in a relatively more hydrophilic section of the pore entrance (Figure 2C&D). We looked at hydropathy of the pore, which assigns hydrophilic amino acids a negative value and hydrophobic amino acids a positive value to examine the overall environment of the pore (42). We found that the inversion from negative to positive hydropathy was located at the LPL active site residues (Figure 2C). At the top of the pore, where we would expect the glycerol backbone is located during glyceride hydrolysis, there is a negative hydropathy score, indicating hydrophilicity. This transitions to a positive score following the active site, suggesting that the hydrophobic fatty acid chain could rest in the remainder of the pore. A 16-carbon fatty acid, such as palmitate, would have a length of ∼24 Å and the ∼30 Å of pore following the LPL active site is more than sufficient to hold that length of hydrophobic molecule. To better understand the ability of acyl chains to fit into the hydrophobic pore we modeled 4 separate ligands into the pore using PyRosetta (43) (Figure 2E, Supplemental Figure 5). The pdb files for these models are available as supplemental materials. Saturated and unsaturated ligands were able to fit in the pore.

### Comparison of active LPL to other open lipase structures

We next compared our structure to other known lipase structures, including structure of pancreatic triacylglycerol lipase (PTL) with the lid peptide in an open position (21, 29, 44) (Figure 3). PTL shares significant structural homology with LPL, as both have lid peptides that regulate access to the active site and swing open during interfacial activation. Two open PTL structures feature hydrophobic ligands bound in the hydrophobic pocket, phosphatidyl choline and tetraethylene glycol monooctyl ether (TGME), respectively (Figure 3F-I). The lid peptides in these 2 PTL structures and our LPL dimer have shifted into an open state, exposing the active site (Figure 3C,G&I). However, a cross-sectional comparison of these proteins with the LPL dimer shows a substantial difference between the size of the previously characterized hydrophobic pocket and the hydrophobic pore seen in our LPL structure (Figure 3B,F&H). The hydrophobic pore is unique to our structure, which we hypothesize could be due to LPL’s response to the AWI during cryo-grid preparation.

**Figure 3.**
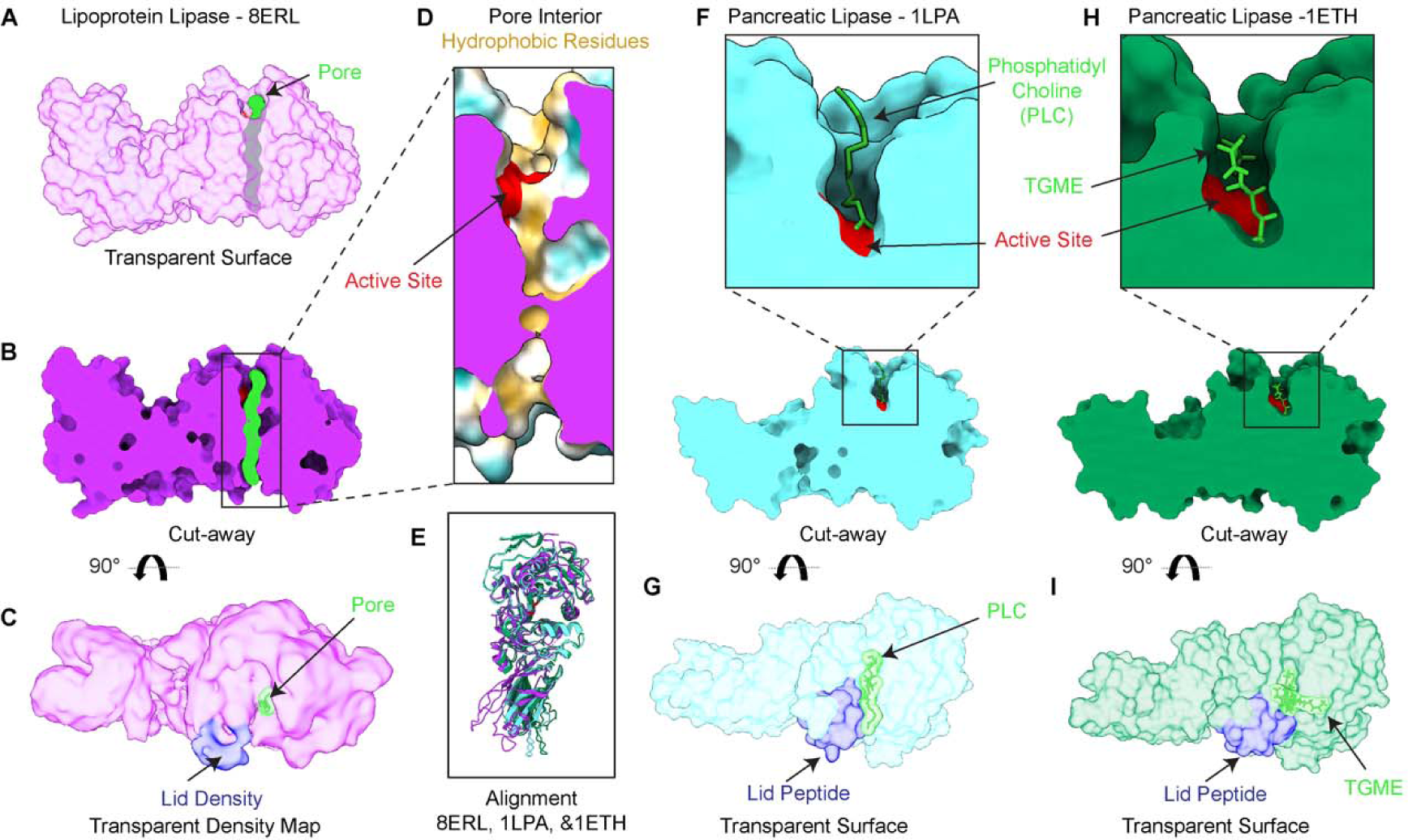
A pore is not present in other open lipase structures. A) A transparent surface representation of a single subunit of LPL (purple) from the LPL dimer, with the path of the pore shown in green. B) A cutaway shows the path of the pore through the protein. C) The density corresponding to the LPL lid domain shown in blue has swung to the side giving access to the active site and pore. D) A closeup look at the LPL pore cutaway, illustrating the active site residues (red) and colored on the surface to show hydrophobic (yellow) and hydrophilic (aqua) residues. The cutaway surface is in purple. E) Alignment of LPL and two pancreatic triacylglyceride lipase (PTL) structures (cyan and green) shows similar folds and well aligned location of the active site residues. F) A cutaway of the structure of human PTL (cyan) aligned to LPL – no pore is observed, only a small pocket adjacent to the active site. The PTL/procolipase complex was crystallized in the presence of phophatidyl choline micelles and bile salt (PDB 1LPA). A zoom-in of the active site (red) in the cross-section shows a diundecyl phosphatidyl choline (PLC) bound. G) PTL in the presence of procolipase was observed with an open lid peptide, revealing access to a hydrophobic pocket, where the PLC bound. H) A cutaway structure of porcine PTL (green) from a PTL/colipase complex, solved in the presence of bile salts and detergents, shows the active site (red) and a tetra ethylene glycol monooctyl ether (TGME) ligand is observed bound in the hydrophobic pocket (PDB 1ETH). I) A transparent surface shows that the lid peptide is in an open state, providing access to the hydrophobic pocket for TGME binding.

### The LPL dimerization interface resembles the GPIHBP1 binding interface and helical LPL interface

We next examined the C-terminal to C-terminal LPL dimerization interface (Figure 4A). Prior structural studies show that LPL has multiple possible interaction interfaces as observed with GPIHBP1 and the LPL helix (Figure 4A-C). Comparison of these interaction interfaces revealed that the novel LPL dimerization interface shared almost complete overlap with the GPIHBP1 binding interface (Figure 4D&E). To further explore this shared binding interface, we analyzed it using PDBePISA (45). The interface involves 14 amino acids from each LPL subunit, and 12 of these residues are shared with the LPL/GPIHBP1 interface (Supplemental Table 1). The homodimerization interface also overlaps with interfaces involved in formation of the inactive LPL helix. The base subunit of the LPL helix is an inactive dimer, which interacts with other inactive LPL dimers through helical interfaces to form a helical filament (5). There was no overlap between the inactive helical dimerization interface and the active homodimer interface (Figure 4G). However, there was overlap with the other helical interfaces, they share 13 interacting residues (Figure 4F). Of these 13 residues, 3 were involved in the C-terminal to C-terminal helical interface, and 11 in the C-terminal to N-terminal helical interface (one residue interacts with both). This suggests that active dimers are incompatible with formation of the repeating helical structure because the helical interaction interfaces are occluded by the presence of the other LPL subunit in the helical interface. All three interfaces – LPL homodimer, LPL/GPIHBP1, LPL helix - share residues Y-G-T-V-A-E (370-375 in bovine LPL and 367-372 in human LPL) and L406, M407, and K409 in bovine LPL (403, 404, and 406 in human LPL). This patch is hydrophobic due to the presence of tyrosine, valine, leucine, alanine, and methionine.

**Figure 4.**
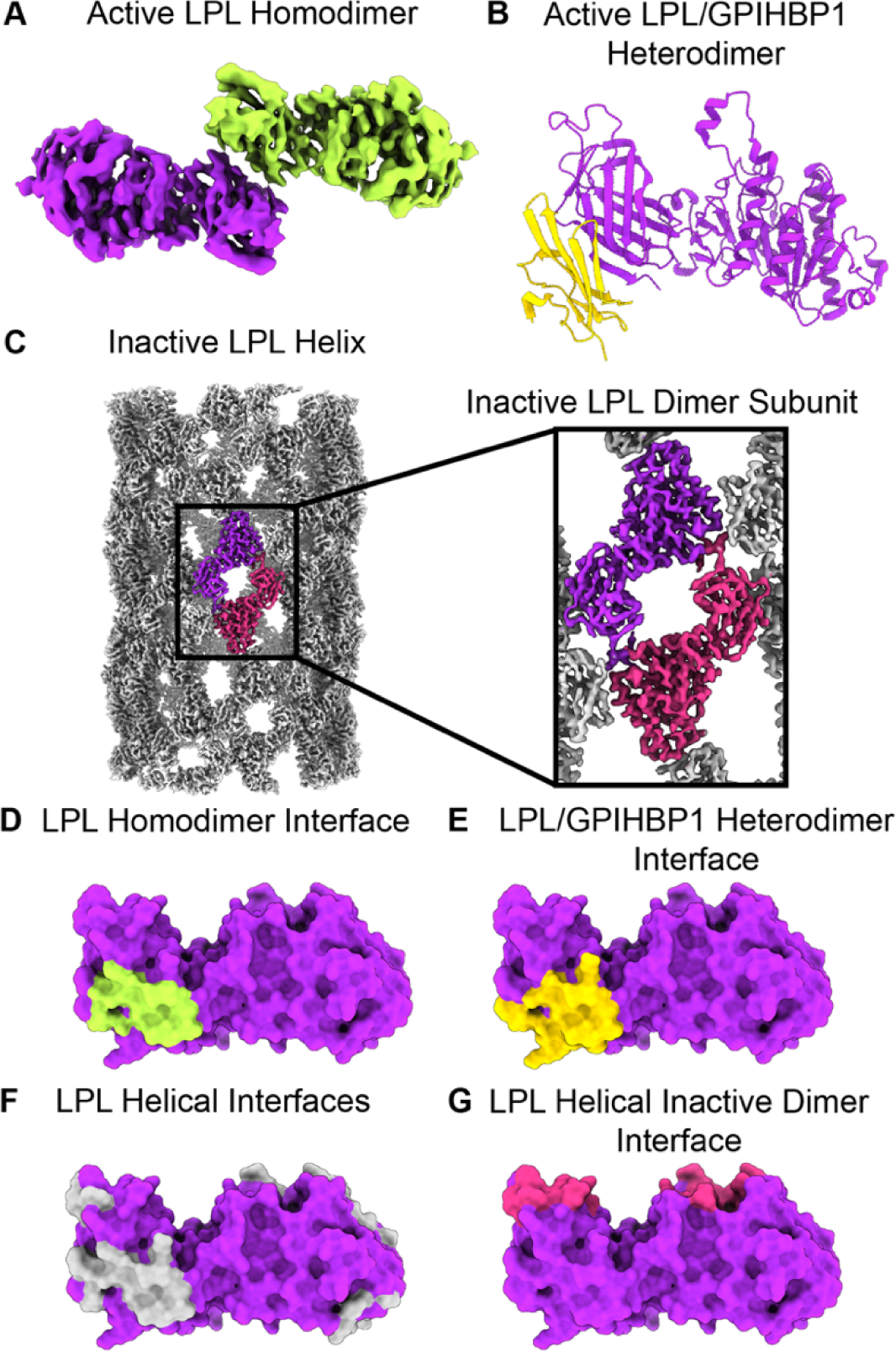
The LPL dimerization interface overlaps with other known interfaces, including LPL/GPIHBP1 and the LPL helix. The structure of LPL (purple) has been solved in 3 distinct states (A) an active homodimer described in this work (one LPL subunit is colored lime green), (B) a heterodimer with GPIHBP1 (gold) (shown as PDB 6OB0), and (C) an inactive helix (EMDB 20673), for which the repeating subunits are inactive LPL dimers (purple and magenta, the rest of the LPL in the helix is colored gray). We used PDBePISA to identify the interacting residues for each of these complexes. D) In the LPL homodimer the interaction interface was located at the C-terminal, and we mapped the residues identified as being buried by the interface onto a space filling representation of one monomer of the homodimer LPL (interface residues, colored lime green). E) The LPL residues interfacing with GPIHBP1 (E, gold) are mapped onto the same LPL, as are the helical interfaces (F, gray) and the inactive dimer interface (G, magenta). The helical interfaces represent the interactions that occur in the LPL helix that are not with the companion LPL subunit found in the inactive dimer. There is an overlap between the active LPL homodimer, LPL/GPIHBP1 heterodimer interface, and LPL helical interfaces.

### LPL oligomeric state is influenced by concentration and additives

We next set out to use an independent method to assess the variability in LPL’s oligomeric state. We did so in part because deoxycholate was needed to prevent LPL helix formation and bile acids are present only at low levels in the blood (∼350 nM of deoxycholate (46)) and display high interindividual variability (47). We wanted to ensure that the higher levels of deoxycholate used to break up helices did not alter the formation of LPL oligomers in solution. First, we used mass photometry to characterize the molecular weight of LPL at low concentrations. Mass photometry is an emerging technique that detects the impacts of protein particles hitting a glass coverslip to determine the molecular weight of the proteins in solution (48). This technique allows for analysis of relatively low concentration protein samples in solution.

At 16 nM LPL, the molecular weight was best fit by a single gaussian distribution (Figure 5A) with a peak at 56 kDa, correlating to a monomer of LPL (50.5 kDa). When the concentration of LPL was increased, a shift toward higher molecular weight was observed (Figure 5A). At higher concentrations (50 and 78 nM) we observe a higher molecular weight shoulder appearing on the monomer peak, suggesting that a second oligomeric species is beginning to form. We fit these concentrations with multiple gaussian distributions, although we do not believe that the molecular weight suggested by the secondary peak (77 and 73 kDa respectively) is an accurate reflection of the exact molecular weight, since it is a subspecies with significantly fewer data points compared to the monomeric peak. However, when we increase the concentration to 100 nM, we see a peak at 99 kDa formed by the majority of the particle impacts, suggesting that at this concentration LPL has transitioned from being primarily monomeric to majority dimeric. At 100 nM we again see a shoulder forming (131 kDa), perhaps indicating that another higher weight oligomer is forming.

**Figure 5.**
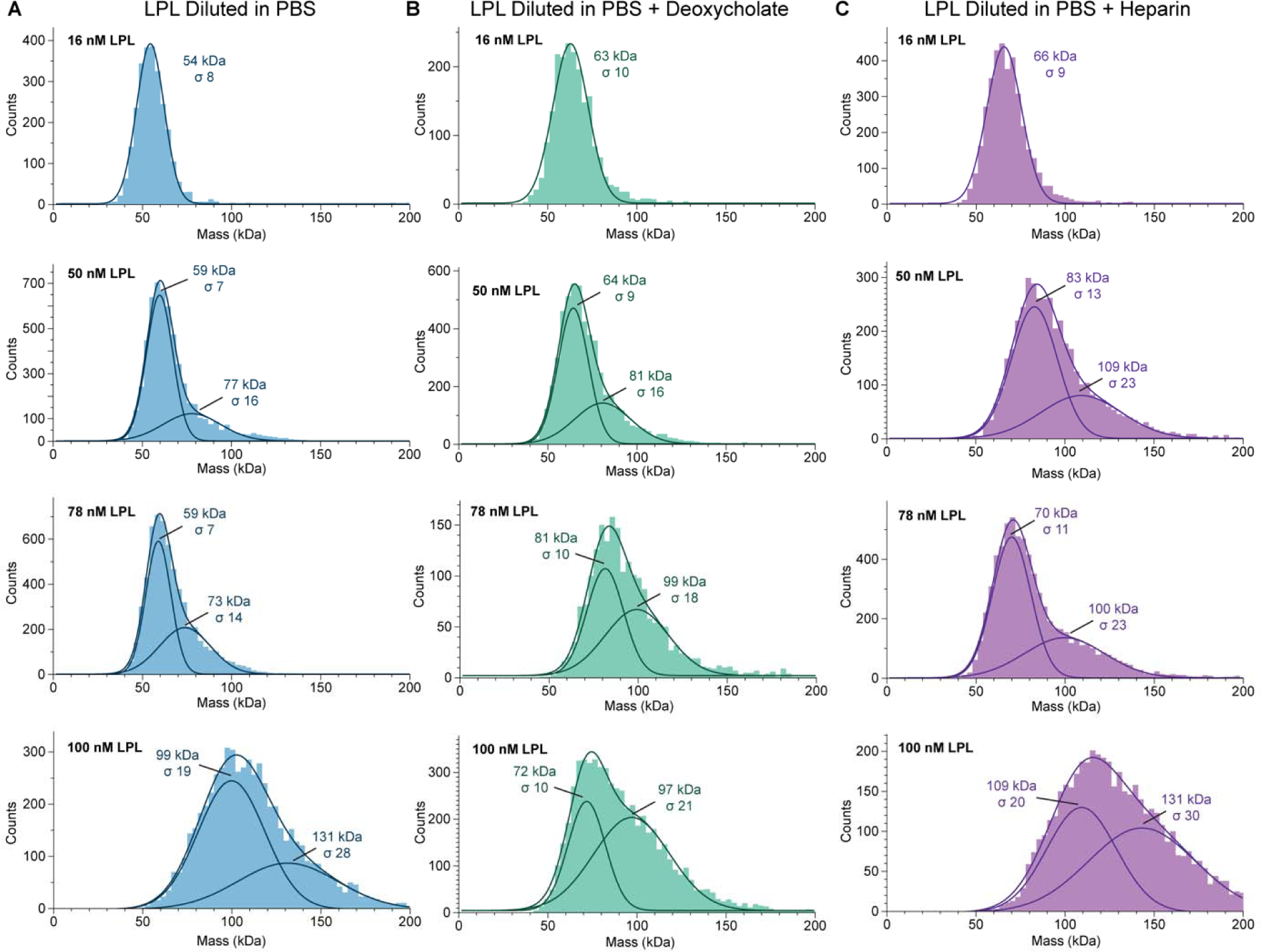
Concentration Drives LPL oligomerization. A) Mass photometry of LPL at increasing concentrations shows that when LPL is analyzed at low concentrations it forms a monomer, and as LPL concentration increases a higher molecular weight shoulder appears. At 100 nM LPL the primary peak corresponds to the molecular weight of an LPL dimer. B) LPL in PBS with 0.25 mM deoxycholate displays a similar trend to LPL without deoxycholate. C) LPL measured in PBS with 27 U/mL heparin also shows the shift to dimerization with increasing concentration. However, the appearance of a dimeric molecular weight peak is observed at lower concentration (50 nM) compared to LPL or LPL with deoxycholate. The data was fitted with single or multiple gaussian distributions to determine an average molecular weight (kDa) and the width of the standard deviation of the fitted gaussian is given by sigma (σ). The combined trace of the multiple gaussians is also show to illustrate the fit of the gaussians to the histograms. LPL monomer theoretical molecular weight = 50.5 kDa.

When LPL at 16 nM LPL was measured in solution containing 0.25 mM deoxycholate, we again saw a single peak of monomeric LPL at 63 kDa (Figure 5B). As we increased LPL concentration in the presence of deoxycholate we saw a similar pattern to LPL alone, with a higher molecular weight shoulder being observed at 50 nM (81 kDa). At 100 nM of LPL with deoxycholate, we still see two peaks, but the primary peak has shifted to a mass indicative of LPL dimerization: 97 kDa. This matches the data for 78 nM LPL with deoxycholate, which also shows two peaks, although with lower occupancy, at 99 kDa. When fit with multiple gaussians, we see there are still contributions from lower molecular weight species at 78 nM and 100 nM (81 kDa and 72 kDa respectively), but as the concentration of LPL increases the ratio of the dimeric protein to the lower molecular weight protein increases, suggesting a correlation between the increasing concentration and the population of LPL dimers.

We also tested an additional LPL additive, heparin, which binds to LPL and has been shown to stabilize higher order oligomers (5). We mixed 27 U/mL mixed molecular weight heparin with LPL and again at 16 nM LPL observed the appearance of a single molecular weight species at 66 kDa (Figure 5C). The LPL monomers observed with deoxycholate and heparin have a slightly higher molecular weight than LPL alone, this may be due to association with heparin or deoxycholate increasing the apparent molecular weight. At higher LPL concentrations of 50 nM and 78 nM, we again observed a shift to multiple gaussian distributions, with minor peaks at 109 and 100 kDa respectively, which correlate to LPL dimers. Interestingly, in the heparin condition the molecular weight matching a dimer appears at a lower concentration than with LPL alone or with deoxycholate. We also see peaks for a lower molecular weight species (83 and 70 kDa respectively). The general distributions at both 50 and 78 nM appear similar, however, at 100 nM we see a complete transition to dimeric oligomers (109 kDa), with further mass for higher order molecular weight species (131 kDa). This data indicates that the oligomerization of LPL is dynamic with respect to concentration, but also suggests that deoxycholate is not significantly affecting the ability of LPL to move through these oligomeric states.

We also confirmed the concentration dependence of oligomer formation seen with bovine LPL is shared by human LPL, by analyzing both 16nM and 50 nM human LPL (Supplemental Figure 6A). At 16 nM human LPL has a primary peak at 58 kDa that primary peak shifts to 99 kDa at 50 nM.

### LPL forms multiple higher order oligomers

Mass photometry is an effective technique for dilute solutions, but we also wanted to capture LPL oligomers from higher concentration solutions and observe any differences in LPL oligomer formation. We used gradient fixation (GraFix) to stabilize oligomers for later analysis (49, 50). We separated both 1 µM LPL and 1 µM LPL with 1 mM deoxycholate using a linear glycerol gradient, which also contained a gradient of glutaraldehyde crosslinker, resulting in individual fractions that contain less than 100 nM of protein. GraFix allows gentle crosslinking, while simultaneously separating the resulting oligomers by size. GraFix is a common method for cryoEM that is not known to induce formation of non-specific oligomers (49). We assayed the resulting crosslinked proteins by both western blot and TAMRA-FP serine-hydrolase active site probe (Supplemental Figure 7). We found that LPL with and without deoxycholate form the same array of oligomers in solution during GraFix (Figure 6A&B). Because crosslinking of LPL oligomers alters their electrophoretic mobility (i.e. a crosslinked monomer of LPL runs at a lower molecular weight than an uncrosslinked LPL monomer), we used mass photometry to confirm the molecular weight of the LPL oligomers observed in the different fractions (Figure 6C&D). Further, by averaging data from multiple fractions, we could confirm the identities of monomers, dimers, trimers, and tetramers of LPL by molecular weight (Figure 6E). We also confirmed that human LPL when crosslinked in solution results in the appearance of the same oligomeric species as bovine LPL (Supplemental Figure 6B).

**Figure 6.**
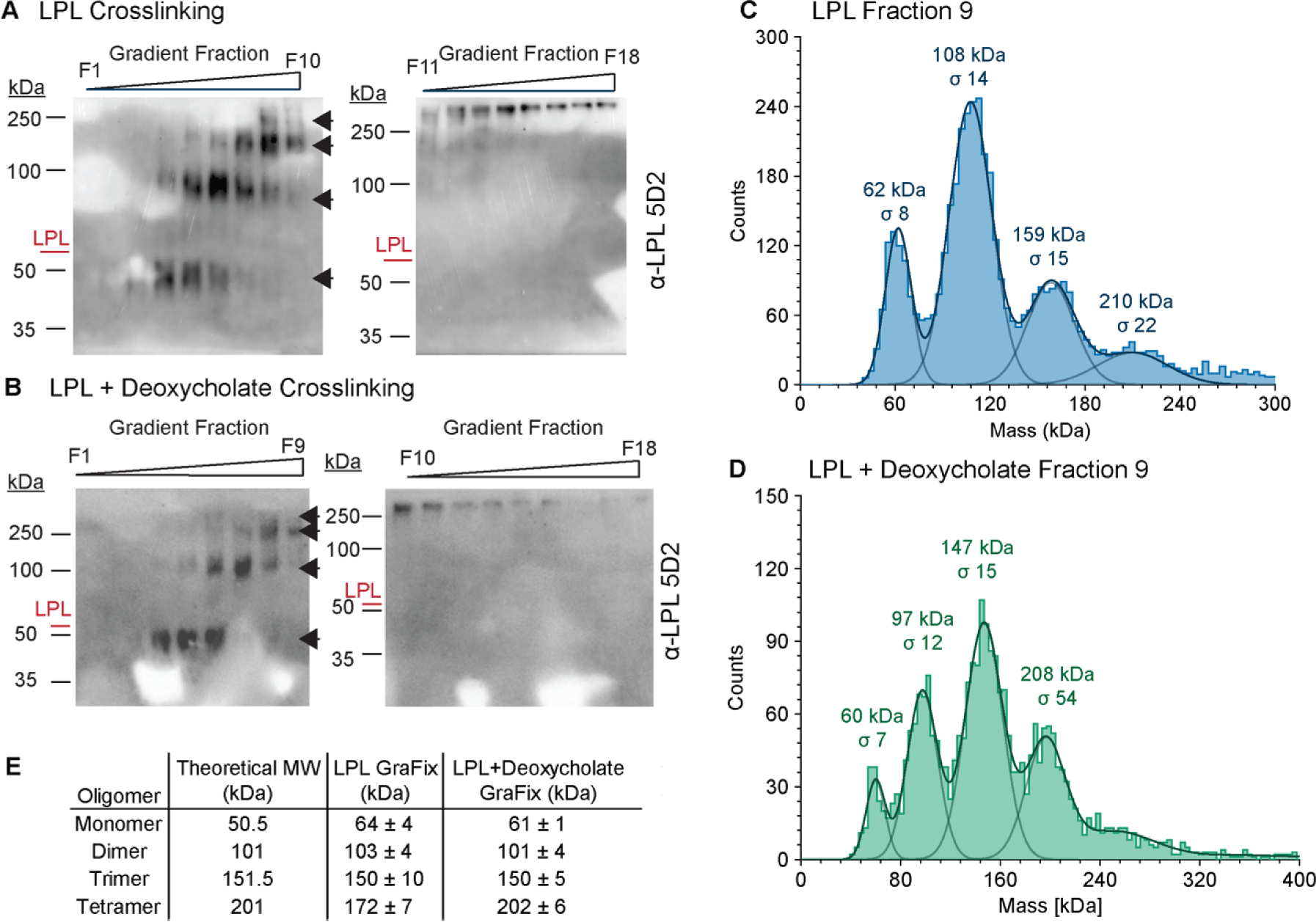
Crosslinking LPL captures monomers, dimers, trimers, and tetramers. To capture and analyze LPL oligomers, we used gradient fixation (GraFix) to gently crosslink and separate protein by molecular weight. LPL alone (A) and with deoxycholate (B) showed an array of oligomers forming. The position of uncrosslinked LPL is marked in red. Blots were developed using the monoclonal LPL antibody 5D2, and the first observable LPL oligomer runs below the uncrosslinked LPL molecular weight – likely due to crosslinking altering its electrophoretic mobility. At least 4 distinct LPL oligomers are observed in both samples (black arrows). The similarity of LPL with and without deoxycholate suggests that the deoxycholate does not alter the overall oligomeric state of the protein. The results of the gradient fractionations in A & B were analyzed using mass photometry to determine the exact molecular weight of their components. C) LPL fraction 9 and D) LPL with deoxycholate fraction 9 are shown as representative data. The data was fitted with multiple gaussian distributions to determine the average molecular weight (kDa) and the width of the standard deviation of the fitted gaussian, sigma (σ). Multiple fractions were analyzed to collect a high quality gaussian distribution of each oligomer. E) Results for peak molecular weight gaussians were averaged and listed in this table as +/- the standard deviation of 3 or more separate measurements. We observe consistent values for monomer, dimer, and trimer for LPL with and without deoxycholate. Molecular weight varied for the likely tetrameric oligomer between the 2 conditions, likely due to fewer data points collected for the LPL tetramer. This difference in distribution can likely be attributed to fractionating by hand and not inherently less tetramer existing in the LPL samples.

In contrast to the solution crosslinking data we did not observe monomers or trimers in our cryoEM data. However, in addition to LPL dimers we did observe some 2D classes that were apparent tetramers of LPL (Supplemental Figure 8A). The tetramer formed when two LPL dimers came together. One LPL subunit of each oligomer appeared to rotate, forming a head-to-tail interaction that may mimic the inactive dimer, which is the repeating subunit of the LPL helix (5) (Supplemental Figure 8B&C).

## Discussion

LPL has long been known to form an active dimer (13, 14, 16–18) and our cryoEM structure reveals the molecular architecture of this complex, including an unexpected C-terminal to C-terminal interaction interface. LPL was previously predicted to form a head-to-tail (N-terminal to C-terminal) dimer (16, 51, 52). This head-to-tail arrangement was observed in the inactive LPL helix and the heterotetramer of the LPL/GPIHBP1 crystal structures. However, in our new structure LPL clearly forms a tail-to-tail dimer. Analysis of the homodimerization interface shows significant overlap with the LPL/GPIHBP1 interface and the LPL helical interfaces. This suggests that LPL homodimerization, binding to GPIHBP1, and LPL helix formation are all mutually exclusive states. Therefore, when LPL is bound to GPIHBP1, it is not able to form a dimer, but upon dissociation from GPIHBP1, LPL would be able to form a homodimer. This C-terminal interaction site shared in all three known LPL structures (5, 7, 8) may play a role in stabilizing LPL, whether in an active or inactive form, enabling structural characterization. The C-terminal to C-terminal arrangement also allows the tryptophan loop, lid peptide, and active site residues of both LPL subunits to interact with substrate at the same time, which was not seen in previous models of the LPL dimer (51).

We also observed a hydrophobic pore adjacent to the LPL active site, formed by rearrangement of the residues lining the pore. The closest analog we found in the literature to this open pore is the tunnel found in C. rugosa lipase, the tunnel gates entrance to the active site (30). However, the C. rugosa lipase has not been shown to have a separate exit as we see in the LPL pore. Our analysis of the LPL pore revealed a pore ideally suited to harbor the hydrophobic fatty acid tail of a triglyceride. The hydrophobic pore could provide an additional layer of substrate discrimination for mammalian lipases beyond the previously identified lid peptide and hydrophobic binding pocket. In that regard, the pore might utilize a similar specificity mechanism to the tunnel seen in C. rugosa lipase and other homologs (53, 54). This could provide an explanation for why LPL preferentially hydrolyzes the sn-1 and sn-3 position on triglycerides, as the sn-2 acyl chain interacts in a separate hydrophobic pocket facilitating enhanced substrate recognition of the sn-1 or sn-3 position by the hydrophobic pore. This is suggested in our PyRosetta model of a triglyceride in the LPL structure (Supplemental Figure 5). Overall, ligand specificity for LPL would then be conferred by a combination of the lid peptide, hydrophobic pocket, and hydrophobic pore.

The pore layout also suggests the existence of a unidirectional mechanism where the FFA product exits from the bottom of the pore, which would allow the FFA to enter the capillary, while the remaining monoacyl-or diacyl-glycerol returned to the chylomicron (Figure 7). This mechanism brings previously known substrate binding regions together and integrates them with the existence of an active site pore. Further, once a triglyceride is hydrolyzed, the diacyl-, monoacyl-glycerol, or glycerol is then able to exit from the entrance of the pore and diffuse back to the surface layer of the TRL (55), while the liberated FFA can pass unidirectionally through the pore and directly into the capillary. This mechanism has the appeal of being very efficient, as LPL would not be required to repeatedly undergo interfacial activation in order to bind to a new ligand or to dissociate from the TRL to release FFA. The processive hydrolysis of ligands by unidirectional travel of the FFA products could also be a mechanism used by other mammalian lipases.

**Figure 7.**
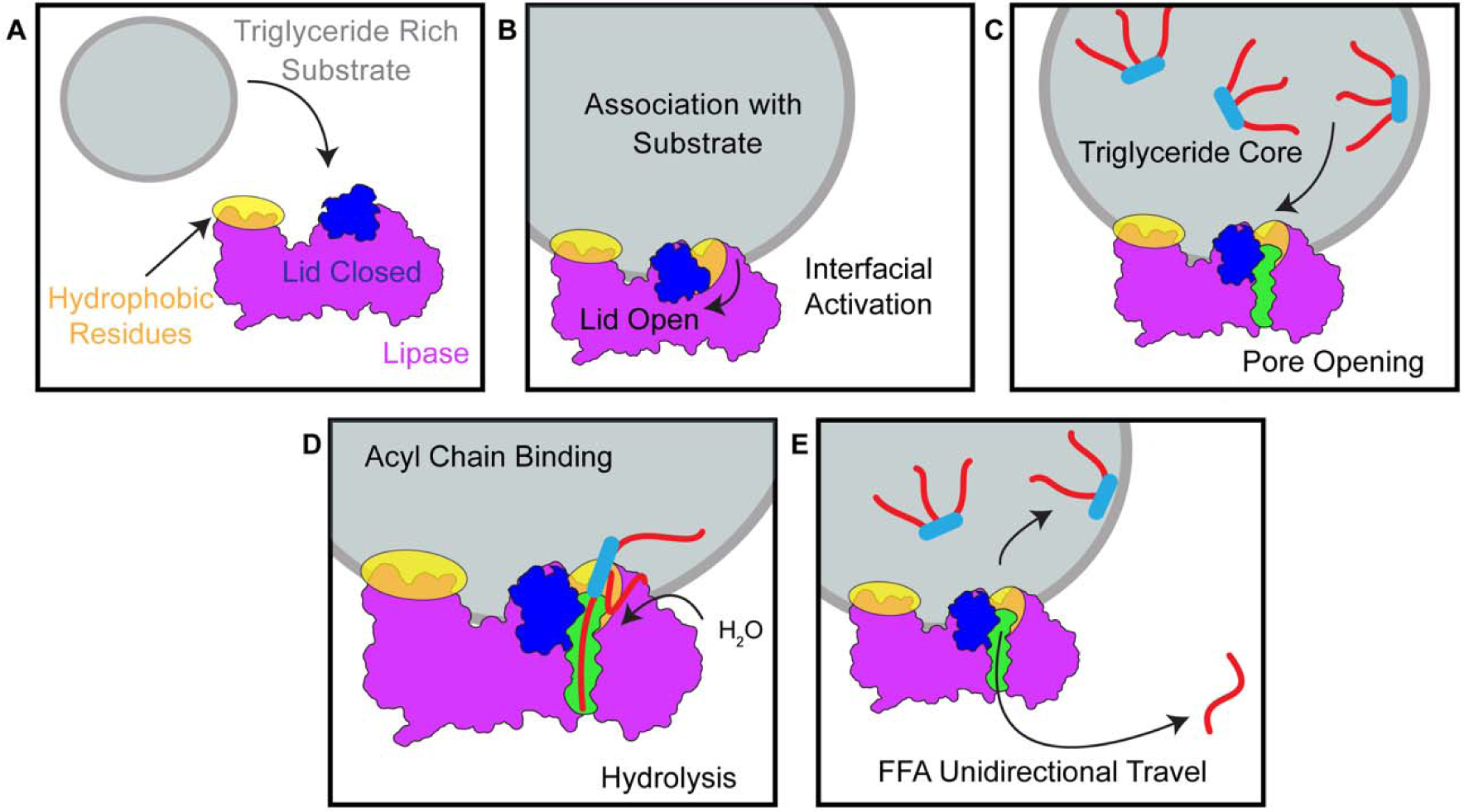
Proposed Unidirectional Model for Triglyceride Hydrolysis by LPL. A) Prior to associating with a substrate, the lid peptide covers access to the active site. Hydrophobic tryptophan residues in the C-terminal domain of LPL are involved in substrate recognition (40). B) After association with a substrate, LPL undergoes interfacial recognition, where the association with a nonpolar-aqueous interface facilitates the lid peptide swinging to the side. C) Following lid-peptide opening, the presence of triglyceride or other hydrophobic ligands trigger formation of an acyl-chain binding pore. D) The hydrophobic acyl chain enters the pore positioned so that the lipase active site residues can hydrolyze the connection of the acyl chain to the backbone. E) Following hydrolysis, the free fatty acid (FFA) exits the pore from the opposite side it entered, resulting in unidirectional FFA travel. The remaining ligand can continue to be hydrolyzed at other acyl chains or transfer back to the hydrophobic substrate. This unidirectional model efficiently allows FFA to transfer away from their original source and be bound by proteins, such as albumin, that can facilitate their travel to surrounding tissues for continued energy processing.

Previous work on the C. rugosa lipase tunnel has shown the mutations in the tunnel can tune the substrate specificity of the lipase (54). Mutations leading to LPL deficiency have been extensively characterized to better understand the causes of familial hyperchylomicronemia. We therefore decided to compare the amino acids that line the pore in LPL to known mutations resulting in LPL deficiency and look at their influence on substrate specificity. We identified R102S in humans (R105 in bovine structure), which leads to LPL deficiency (56, 57), and L279R in humans (L282 in bovine structure), which leads to lower LPL specific activity (58). These two mutations which are both known to have deleterious effects on human LPL activity localize to the pore exit (Figure 2D). (In literature predating structures of LPL, these residues were referred to as R75 and L225 in humans, accounting for removal of the signal sequence. We will refer to these residues by their numbering in recent LPL structures for consistency.) The LPL R102S variant was identified as one of two mutations in a patient with compound heterozygous loss of LPL function resulting in severe chylomicronemia (57). The R102S mutation results in loss of positive charge at the pore exit (Figure 2D and Supplemental Figure 9A&C). In previous work LPL with the R102S mutation was expressed in and purified from cultured cells, and a reduced amount of active enzyme was produced relative to the WT LPL. Interestingly, when corrected for protein concentration, the WT LPL showed higher specific activity on a long chain, triolein substrate relative to LPL R102S, but LPL R102S showed higher specific activity on the small soluble substrate, para-nitrophenyl butyrate (57). The LPL L279R mutation substitutes a hydrophobic leucine residue with a bulky, positively charged arginine residue. Residue 279 is positioned directly inside of the pore close to the exit (Figure 2D). Homology modelling in our structure suggests that this bulkier arginine residue may block the pore (Supplemental Figure 9B&D). A patient heterozygous for the L279R mutation developed severe chylomicronemia during pregnancy (58). In vitro studies revealed that arginine or proline substitutions at LPL residue 279 resulted in production of 30% or 70% of WT protein, respectively. However, both LPL variants had no specific activity with a long chain triglyceride substrate, specifically a triolein emulsion (58).

Thus, both LPL pore mutants identified from chylomicronemic patients could fold and be secreted. However, both variants showed a specific defect on long chain triglyceride substrates, suggesting the pore observed in our structure has physiological relevance for hydrolysis of specifically long chain ligands, which would fit in the hydrophobic pore. Whether these mutations affect release of LPL product or alter substrate specificity requires further investigation.

The LPL dimer structure was solved in the presence of deoxycholate to create a homogenous sample. However, the LPL dimer does not form solely because of deoxycholate treatment. The appearance of the LPL dimer is linked to increasing LPL concentration. We used crosslinking to capture LPL dimers in samples both with and without deoxycholate. We observe that the distribution of oligomers captured in both solutions are similar, with monomers, dimers, trimers, and tetramers observed by SDS-PAGE and mass photometry. It has been previously reported that other lipases are activated or stabilized in the presence of bile salts, for example bile salt dependent lipase (59, 60). Deoxycholate may act in a similar fashion to stabilize LPL and it might play a role in shifting the lid peptide to an open position, allowing access to the active site (61). Indeed, we see that the density corresponding to the lid residues in our map are shifted away from the opening of the active site. This is a different orientation for the lid peptide than was observed when LPL was crystalized in the presence of an active site inhibitor (7). Given that deoxycholate can serve as a natural activator, this position of the lid, where it has swung to the side, likely reflects the physiological movement triggered by interfacial activation that allows triglycerides to enter the active site. Indeed, there is significant structural overlap between the open lid density in LPL and the open lid peptides in PTL structures (Figure 3).

LPL had a pronounced preferred orientation in our data, which likely resulted from its interaction with the AWI (37). We speculate that during grid freezing, LPL interacted with the AWI as though it were a hydrophobic substrate. The interface between the aqueous buffer and air thus mimicked the nonpolar to aqueous interface that triggers interfacial activation of LPL in vivo. Accordingly, the AWI pseudo-substrate favored formation of active dimers of LPL primed for substrate hydrolysis (13, 14) with the lid-domain shifted to be in an open state and the active site pore exposed. We speculate that other lipases may also adopt a preferred orientation at the AWI in cryoEM. The tilting and particle picking strategy described here may be useful for solving structures of other lipases in active states possibly with open hydrophobic pores.

In our cryoEM data we identified classes corresponding to a dimer and tetramer of LPL – but not the monomer or trimer, which we observed only in solution by mass photometry and crosslinking. We believe the AWI pseudo-substrate qualities may explain this discrepancy. It has previously been observed that LPL’s oligomeric state is sensitive to the presence of an aqueous-nonpolar interface (5). In an aqueous solution, such as the one where we performed our crosslinking experiments, LPL was not exposed to a potential substrate interface. This allowed LPL to exist in a variety of oligomeric states, including ones that may be inactive, such as the monomer and trimer. However, during cryo-grid preparation, exposure to the AWI triggered LPL to adopt its active state, leading to formation of active dimers. As for the tetramer, it would likely be partially active as 2 of the LPL subunits are oriented to the AWI (Supplemental Figure 8). The other 2 subunits share similarity with the inactive dimer found in the LPL helix, which we suspect may be a result of the high concentration of LPL used to freeze grids. At high concentrations LPL favors helix formation to efficiently pack and store LPL without risk of inactivation (5), but the presence of active LPL dimers due to the AWI and deoxycholate precluded helix formation in this case.

Our structure shows that dimeric LPL exists in vitro, leading to the question of where dimeric LPL might exist in vivo. Originally, when dimeric LPL was identified as the active form of LPL, it was thought that dimeric LPL (17) was attached to HSPGs in the capillaries (62) to interact with lipoproteins. The discovery of the LPL/GPIHBP1 interaction (6), and the fact that LPL forms a heterodimer with it (7, 8), changed the conventional thinking on the matter. Previous work has shown significant association of LPL and lipoproteins in post-heparin plasma (heparin dissociates LPL from GPIHBP1) (10, 32) and illustrates that LPL does not require GPIHBP1 to associate with a lipoprotein (40). Nor indeed is GPIHBP1 required for LPL to hydrolyze lipoprotein triglycerides, as free LPL can readily hydrolyze lipoproteins in plasma samples (10, 32). To determine which lipoproteins LPL primarily associates with, plasma samples were treated with a lipase inhibitor immediately following collection, and LPL was found predominantly on VLDL particles (10). By contrast, when an inhibitor was not used, LPL was found on cholesterol-rich lipoproteins, such as high- and low-density lipoproteins (HDL and LDL) (18). These data show that LPL remains active on lipoproteins after removal from GPIHBP1 and continues to hydrolyze triglycerides until they are depleted. It has also been shown that in pre-heparin plasma (i.e. untreated plasma) LPL still associates with lipoproteins, specifically VLDL (10). Recent work using fluorescence to monitor the LPL/GPIHBP1 complex has shown that LPL dissociates from GPIHBP1 in the presence of its substrate, chylomicrons, but also when excess product, free fatty acids (FFA), are added (63). The release of FFA from chylomicrons by LPL may promote dissociation of the LPL/GPIHBP1 complex (63).

Beyond hydrolyzing triglycerides, LPL binds to lipoprotein remnants and facilitates their uptake by the liver (12). Prior experiments using sandwich enzyme-linked immunosorbent assays (ELISAs) showed that dimeric LPL is responsible for mediating lipoprotein binding to the low-density lipoprotein (LDL) receptor-related protein (LRP) (64, 65). Heparin does not affect the association of LPL with lipoproteins but does disrupt LPL interaction with GPIHBP1 and LRP (66). We hypothesize that following dissociation from GPIHBP1, in the presence of substrate, LPL would favor formation of an active dimer (Figure 8). This would stabilize LPL and allow it to hydrolyze triglycerides from the core of the lipoprotein until they were depleted. LPL would then remain in its dimeric state to facilitate remnant uptake by the liver (64). The transition of a chylomicron to a chylomicron remnant entails a reduction in the circumference of the lipoprotein, as triglycerides are hydrolyzed, potentially the resulting alteration in lipoprotein curvature or exposure of new residues from chylomicron constituent proteins could serve as a trigger for LPL dissociation from GPIHBP1 in addition to production of FFA.

**Figure 8.**
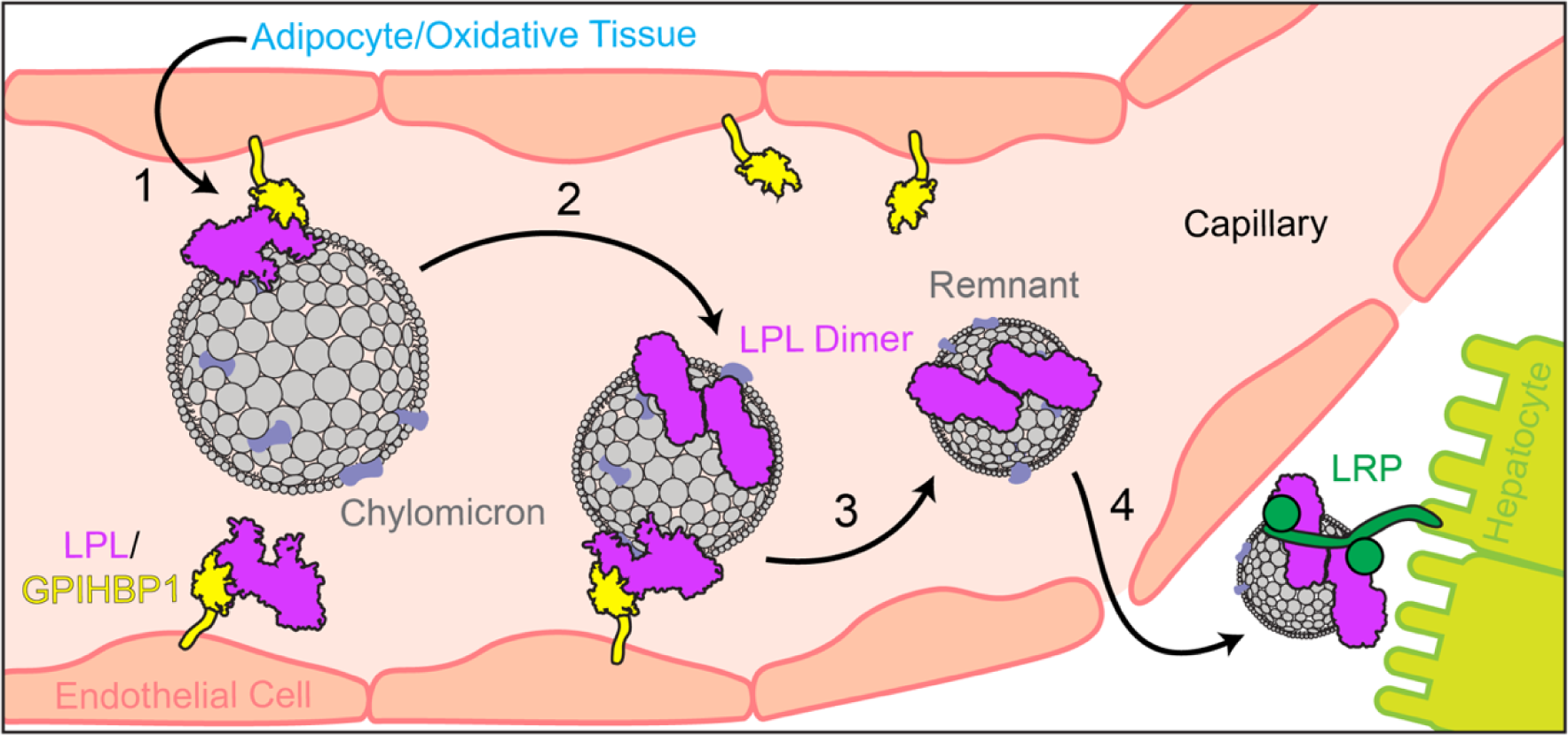
Proposed model of LPL dimer formation in the capillary. LPL is synthesized in the adipose and oxidative tissues and secreted into the interstitial space, where it binds to GPIHBP1 as a heterodimer (7, 8). 1) GPIHBP1 transports LPL across the endothelium into the capillary (6), where it interacts with triglyceride rich lipoproteins, including chylomicrons and VLDL (9). 2) LPL hydrolyzes triglycerides to release free fatty acids and can dissociate from GPIHBP1 (63) – potentially forming a LPL homodimer to maintain hydrolysis activity and protein stability on the lipoprotein particle. 3) Following continued triglyceride hydrolysis, the lipoprotein remnant is released from all GPIHBP1 associations (63) keeping it tethered to the capillary wall - LPL homodimers remain on the remnant (10). 4) The lipoprotein remnant and attached LPL homodimer travels through the capillaries to the liver and pass into the space of Disse, where the remnant binds to hepatic transmembrane protein, LRP. LRP has previously been shown to interact with LPL homodimers to enhance uptake of low-density lipoprotein remnants into hepatocytes (64, 65).

The potential for the unidirectional mechanism for mammalian lipases to facilitate transfer of FFA during hydrolysis provides valuable insight into how lipase mutations can lead to disease. It also opens new arenas of analysis for other mammalian lipases given their significant structural and mechanistic similarity to LPL. It is clear from this work and other recent structures that LPL can adopt a diverse series of oligomeric states. LPL, and lipases in general, play crucial roles in providing energy to the body, but it is equally important that they are active only in the correct milieus. Changes to quaternary structure may allow LPL to self-regulate as it navigates its different roles moving from an inactive form stored in adipocyte vesicles (5), to a heterodimer on the capillary wall (7, 8), and finally onto lipoproteins as a homodimer to eventually facilitate uptake by the liver (64).

## Materials and Methods

### bLPL Purification

Bovine LPL was purified from raw milk using heparin sepharose chromatography as previously described (5).

### Lipase Activity Assay

LPL activity was assayed using an enzyme coupled fluorescence assay to detect non-esterified fatty acids (NEFA) with VLDL as the substrate. Concentrated LPL (∼1 mg/mL) was dialyzed overnight at 4°C in 500 mM NaCl, 20 mM Tris-HCl pH 8 using a 3.5 kDa slide-a-lyzer. When indicated, 1 mM sodium deoxycholate was added to LPL prior to dialysis. A final concentration of 10 nM LPL was used in assays and the starting concentration of the dialyzed LPL was determined by nanodrop.

This assay was performed as described previously (34). Immediately prior to preparing the assay, LPL was diluted to 50 nM in phosphate buffered saline (PBS) with fetal bovine serum (FBS). The final concentration in the well was 10 nM LPL, 0.2x PBS, and 2% FBS. The reaction was started by the addition of the enzymes needed to detect NEFA, VLDL (Athens Research & Technology, 12-16-221204), and Amplex UltraRed (ThermoFisher, A36006). The final concentrations in the well from this mixture were: 133 mM KPO_4_ pH 7.5, 150 mM NaCl, 3.3 mM MgCl_2_, 4.4 mM adenosine triphosphate, 1 mM Coenzyme A (CoA), 0.1 U/mL Acetyl-CoA synthetase (ACS), 6 U/mL horseradish peroxidase, 5 U/mL Acetyl-CoA oxidase (ACO), 0.2 mg/mL fatty-acid free bovine serum albumin (BSA), 0.05 mM Amplex UltraRed, and 200 μ<u>g</u>/mL triglycerides in human VLDL. Assays were conducted in a black-walled, 96 well plate. Immediately following addition of the substrate, fluorescence was monitored using a M5 Spectramax plate reader at 37°C, excitation at 529 nm and emission at 600 nm, with a 590 nm cut-off filter. Initial rate was determined from the first 180 sec of the reaction. 3 biological replicates were conducted for each assay.

### CryoGrid Preparation

LPL at a concentration 1 mg/mL or higher was mixed with 1 mM sodium deoxycholate and dialyzed into 500 mM NaCl, 20 mM Tris-HCl pH 8 at 4°C either with or without 1 mM deoxycholate in the dialysis buffer. Samples were dialyzed overnight or for a minimum of 5 hours. Following dialysis, samples were spun to remove potential aggregates and transferred to a new tube. The concentration of the resulting LPL was verified using a NanoDrop. Several additives were tested to reduce LPL’s preferred orientation, but none were successful in altering the preferred orientation. Samples were vitrified using a Vitrobot set at 4°C and 100% humidity. UltrAuFoil R1.2/1.3 and R0.6/1 300 mesh grids were used. Grids were cleaned using a Tergeo Plasma Cleaner (PieScientific) with settings for direct plasma at 15 W with a 255 duty ratio, in an atmosphere with 2.5% oxygen and 7.5% argon for 1 minute. 3 µL of each sample was applied to the grid, pre-blotted for 10 sec, blotted for 3-4 sec, and plunge frozen in a liquid ethane/propane mix at −180°C.

### CryoEM Data Collection

Grids were imaged using a 200 kV Talos Arctica equipped with a Gatan K3 camera. The microscope was operated using SerialEM (67) and data were collected using beam-image shift with a 5×5 grid to speed data collection (68). Beam-tilt compensation was calibrated in SerialEM to reduce the residual phase error from large beam-image shifts. Data was collected over 2 sessions from two different grids (UltrAuFoil R0.6/1 and R1.2/1.3) prepared with 0.55 mg/mL of LPL that had been dialyzed into 1 mM sodium deoxycholate in 500 mM NaCl and 20 mM Tris-HCl pH 8. All data sets were collected at a nominal magnification of 45,000x (0.89 Å/pixel) with a total flux of 55.1 e^-^/Å^2^. In total 13,748 movies were collected at 0° (4659), 30° (3275), and 45° (5814) tilts.

### CryoEM Data Processing

All data were processed using the Cryo-EM Single Particle Ab-Initio Reconstruction and Classification (cryoSPARC) software suite (39). Raw frames were aligned using patch motion correction with dose weighting followed by patch contrast transfer function (CTF) estimation. The data taken for each tilt (0°, 30°, 45°) were kept separate for pre-processing and particle picking. Micrographs with low resolution CTF fits and high total full-frame motion distance were removed leaving 3566 micrographs at 0° tilt, 2135 at 30° tilt, 4525 at 45° tilt.

Initial processing yielded a low-resolution density map, which was highly anisotropic due to a preferred orientation of LPL on the grid. This map was filtered to 15 Å and used to create templates for particle picking and 2D classification. 2D classes with visible secondary structure were selected and filtered for the best 10% of particles per class using the cryoSPARC class probability filter. The remaining 90% of particles from selected classes and the particles from classes that were not selected were combined and subjected to another round of 2D classification. Following multiple rounds of 2D classification, the highest probability particles from the best classes for each round of classification were used to train the Topaz neural network (38) using the cryoSPARC wrapper. Topaz-picked particles were 2D classified and any rare views (non-preferred orientation) were selected, filtered to select the particles that best fit the non-preferred orientation 2D classes, and fed back to Topaz for another round of training. This process was repeated iteratively until no additional rare view particles were found by Topaz picking.

At this point, particles corresponding to rare views were recombined with the preferred orientation particles found in the initial Topaz picking and checked for duplicates. After generating a complete particle set for each tilt using the above method, the unbinned particles for all tilts were extracted using a 320-pixel box size and combined, resulting in a combined dataset of 1,035,101 particles. Ab-initio model generation was used to find a subset of particles with reduced anisotropy. This was followed by heterogenous refinement and homogenous refinement with C1 symmetry. We then applied C2 symmetry to the homogenous refinement and took the best resulting model with C2 symmetry, which included 527,205 particles. The result of homogenous refinement was then subjected to non-uniform refinement (69) minimizing over a per-particle scale. A final map with a GSFSC resolution of 3.9 Å was obtained. Local and global CTF refinement did not improve the map, possibly due to the low molecular weight of the target protein.

The final map was submitted to the 3DFSC server (36), showing a GSFSC resolution of 3.91 Å and a sphericity of 0.85 out of 1. A sphericity of 1 would indicate that the particles that went into making the map sampled every orientation evenly. This indicated that our tilting strategy was able to overcome the preferred orientation of our sample (36). Analysis of the model found that the hand of the alpha helices was flipped, so the hand was flipped with cryoSPARC volume tools. The final map was sharpened with a map-sharpening B-factor of −100 Å^2^. The distribution of particles to the final map was calculated using UCSF PyEM v0.5 (70).

### Model Building

The structure of an LPL monomer from the inactive LPL helix (PDB 6U7M) was fit into the sharpened map (5) using PHENIX (71, 72). This was followed by iterative rounds of PHENIX real-space refinement and manual adjustment in Coot (73). C2 symmetry was applied to the model and used to generate the other LPL subunit. One round of PHENIX Rosetta Refine (74) was performed on the LPL dimer, followed by further iterative rounds of PHENIX real-space refinement and adjustment in Coot. The final model has a cross correlation efficient of 0.77 with the map; additional statistics can be seen in Table 1.

### Structure Analysis

Structures were visualized using ChimeraX (75). To characterize the LPL active site pore, we used MOLEonline (41). Initial identification of the pore was done with the bottleneck radius parameter set to 1.2 Å with a tolerance of 3. Residues at the top and bottom of the pore were selected as the starting and end points. Overlay of the LPL dimer with other lipase structures was performed in ChimeraX with Matchmaker (75).

### Fatty Acid Substrate Modeling

Models for ligands modeled into the LPL pore were obtained from the PDB: palmitate (PLM), oleate (OLA), linoleate (EIC), and triglyceride (TGL). Ligands were fit into the pore manually, overlaying them with the pore representation from MOLEonline (41) in ChimeraX (75). Coot (73) was used to relax the ligands to reduce clashes. These coot relaxed models were then used to start a PyRosetta (43) minimization run, which allowed both the ligand and LPL to shift in order to generate an improved fit. The PyRosetta modeling resulted in lower repulsion scores for all ligands located in the hydrophobic pore. The models used to start PyRosetta and the final relaxed models are available in the supplemental files.

### Mass Photometry

Mass photometry was performed with a Refeyn Two MP, which uses the impacts of particles hitting a glass coverslip to determine the molecular weight of the particles (48). Coverslips were sonicated in isopropanol prior to affixing 3 mm x 1 mm plastic gaskets (Grace Bio-Labs). BSA was diluted into PBS to test the calibration of the mass photometer before each use – confirming a mass of 66 and 133 kDa, for monomer and dimer BSA respectively. BSA was diluted directly into a single gasket containing PBS. The PBS in the gasket prior to addition of protein was used to focus the mass photometer on the glass-buffer interface. Alternatively, new mass calibrations were created with BSA, Urease, and IgM. A stock of 6.25 µM LPL was diluted into PBS or PBS that been pre-mixed with 110 U/mL heparin or 1 mM deoxycholate. LPL was then diluted into a single gasket containing pure PBS at a 1:4 ratio to collect data. The final concentrations of additives were, therefore, 27.5 U/mL heparin and 0.25 mM deoxycholate. Particle impact events were analyzed as ratiometric contrast data and converted to molecular weight using calibrations with known molecular weight standards (76). Data for each sample was binned in 3 kDa increments and then analyzed with Origin graphing software to fit single or multiple gaussian peaks.

Crosslinked samples from GraFix (described below) were similarly diluted directly into a single gasket containing PBS. Concentration of crosslinked samples were below 100 nM for each fraction. We analyzed multiple GraFix fractions to determine the mass of each observed oligomer.

### Gradient Fixation (GraFix)

Linear gradients from 5% to 20% glycerol were generated in tubes for an SW-60 rotor. 0.2% glutaraldehyde was added to the 20% glycerol to additionally create a linear glutaraldehyde gradient from 0 to 0.2%. The gradients were cooled to 4°C and 150 µL LPL with or without deoxycholate was layered on top of the gradient. The gradients were spun at 38,000 RPM (which has an average force of 148,000 xg) for 18 hours at 4°C in an ultracentrifuge. The gradients were removed from the centrifuge and fractionated from the top in 150 µL fractions. The resulting fractions were analyzed by western blot. The protein oligomers were separated by 8% SDS-PAGE, transferred to 0.22 μm PVDF membrane, and blocked with 5% non-fat milk in TBS-T (20 mM Tris-HCl pH 7.6, 150 mM NaCl, and 0.01% Tween 20). LPL was probed with a monoclonal 5D2 LPL antibody (Abcam, ab93898) at a 1:250 dilution and protein was detected with a HRP-conjugated goat anti-mouse antibody (Southern Biotech) using a 1:5000 dilution. Western blots were developed as previously described (77).

Interestingly, another monoclonal LPL antibody, 4-1a (Millipore, MABS1270), which binds to the N-terminal fragment of LPL, did not detect crosslinked LPL on these blots. To improve our detection resolution, we also performed gradients where we added a final concentration of 1.3 µM TAMRA-FP Serine-Hydrolase probe (TAMRA) (ThermoFisher) to the starting 1 µM LPL mixture. This allowed us to analyze the SDS-PAGE gels without western blotting, using a GE Typhoon scanning at 532 nm. We did not observe any significant differences between the results for western blotting and TAMRA labeling, indicating that the TAMRA probe did not change the oligomeric states adopted by the LPL (Supplemental Figure 3). The TAMRA labeled samples were used for mass photometry – enabling selection of gradient fractions on the same day as gradient fractionation.

### Human LPL

Human LPL samples were analyzed with mass photometry using the same protocol as bovine LPL. A furin resistant LPL mutant was utilized in these experiments, as larger amounts of uncleaved LPL can be isolated from tissue culture with this mutant (78). Crosslinking of human LPL (and comparison bovine LPL) was performed in a 500 nM solution of LPL in PBS with 5% glycerol, 5 mM bis(sulfosuccinimidyl)suberate (BS3) (Thermo, A39266), and 1 µM TAMRA for 30 min at RT. Crosslinking results were analyzed by SDS-PAGE using 8% acrylamide gels and then imaged using tamara fluorescence and western blotting as described above.

### Data availability

The data that support this study are available from the corresponding author upon reasonable request. The Cryo-EM density map was deposited in the Electron Microscopy Data Bank (EMDB) under accession number 28554. Model coordinates have been deposited in the Protein Data Bank (PDB) under accession number 8ERL. Other structures used in this study were obtained from the PDB with accession codes 6U7M (LPL helix), PDB 6OB0 (LPL with GPIHBP1), 8ERL (LPL subunit from the homodimer), 1LPA (PTL/procolipase complex), and 1ETH (porcine PTL). Models for LPL/ligand complexes are available as supplemental files, including before and after minimization with PyRosetta. Raw data for mass photometry is provided as a supplemental excel file.

## Acknowledgments

We thank members of the Neher Lab for discussions and assistance. We thank Dr. Joshua Strauss and Jared Peck at the UNC CryoEM Core and Dr. Nathan Nicely at the UNC Macromolecular Crystallography Core for their assistance. We thank Dr. Brian Kuhlman for assistance with PyRosetta. We thank Dr. Rick Baker and Robert Risti for critical reading of the manuscript. The authors acknowledge support from the National Institutes of Health (R01-HL125654 to S.B.N. and 1K99-GM146024 to K.H.G.) and the American Heart Association (Postdoctoral Fellowship 900354 to K.H.G.).

---

**Supplemental Figure 1:**
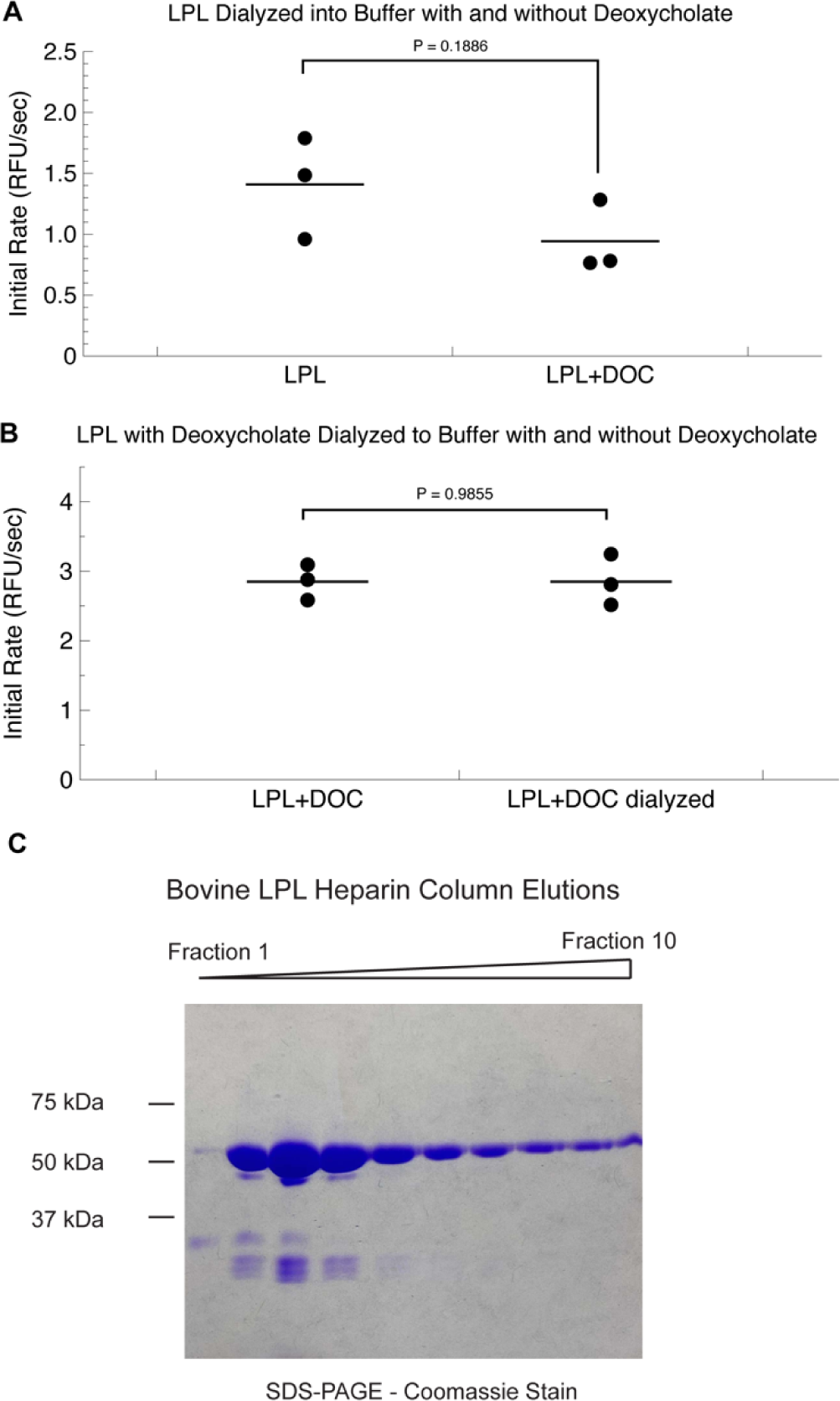
A) LPL activity is slightly reduced following dialysis into buffer containing deoxycholate (DOC) compared to a buffer without deoxycholate. B) LPL incubated with deoxycholate and then dialyzed into a buffer with or without deoxycholate – to remove excess deoxycholate from the protein sample – have virtually identical activity. The slight change in LPL activity seen with deoxycholate is not a result of excess deoxycholate in solution. Assays were performed with 10 nM LPL and 200 μ<u>g</u>/mL triglycerides with VLDL in biological triplicate. C) Coomassie stained SDS-PAGE of bovine LPL purification showing the purity of the preparation.

**Supplemental Figure 2:**
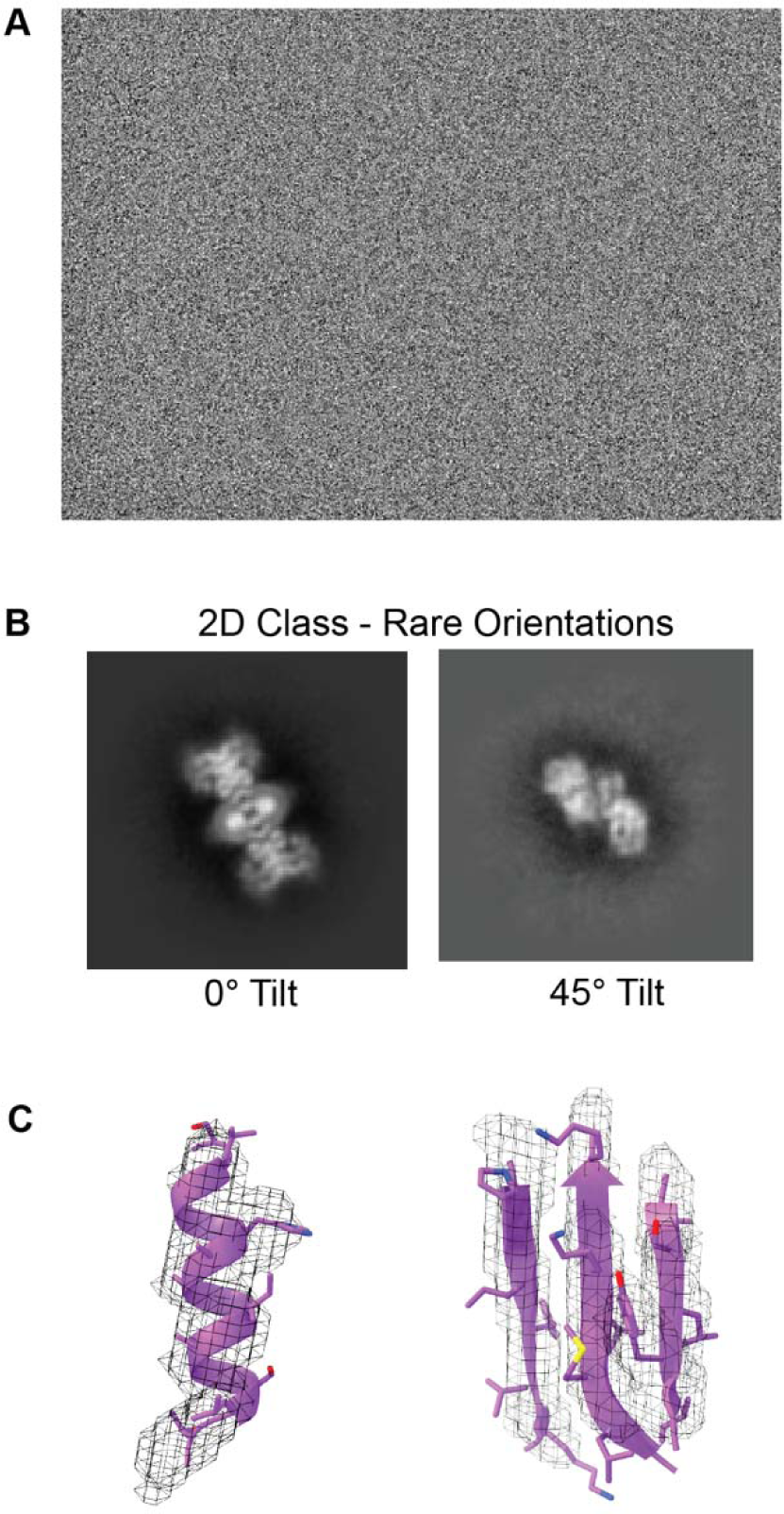
A) Representative micrograph for single particle LPL data. B) Rare 2D classes at each tilt were found using Topaz picking - and combined with the preferred orientation particles to solve the structure. C) The density map had clear separation between the beta-sheets and the turns of the alpha-helices are visible, as can be seen from fitting a portion of the molecular model (purple) into the density displayed as a mesh.

**Supplemental Figure 3:**
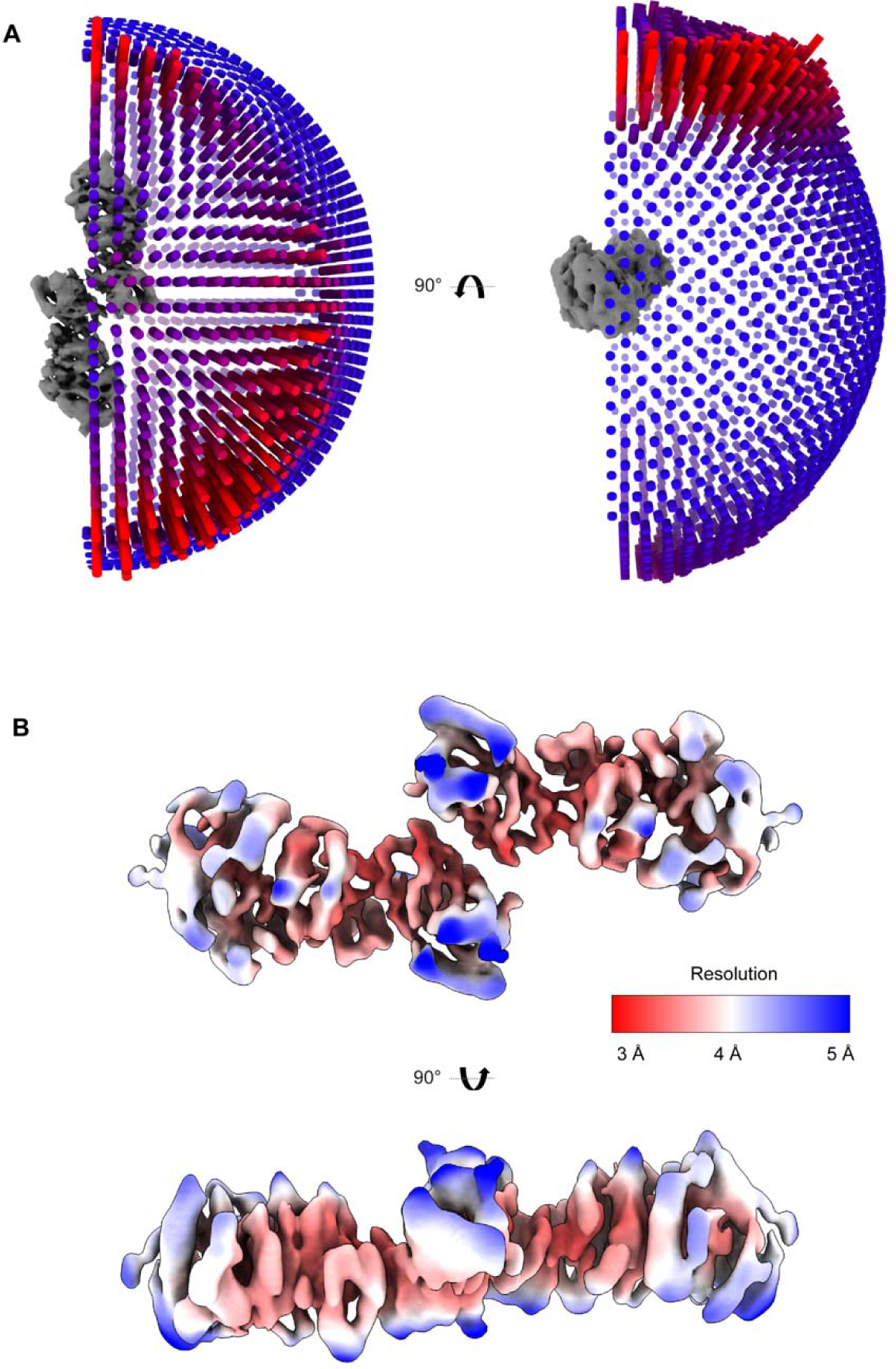
A) Distribution of particles compared to the resulting 3D structure of LPL. The height and color (taller and redder indicate more particles) at each spot indicates the number of particles in the final model that came from a particular angular view. B) A local resolution map of the LPL dimer structure.

**Supplemental Figure 4:**
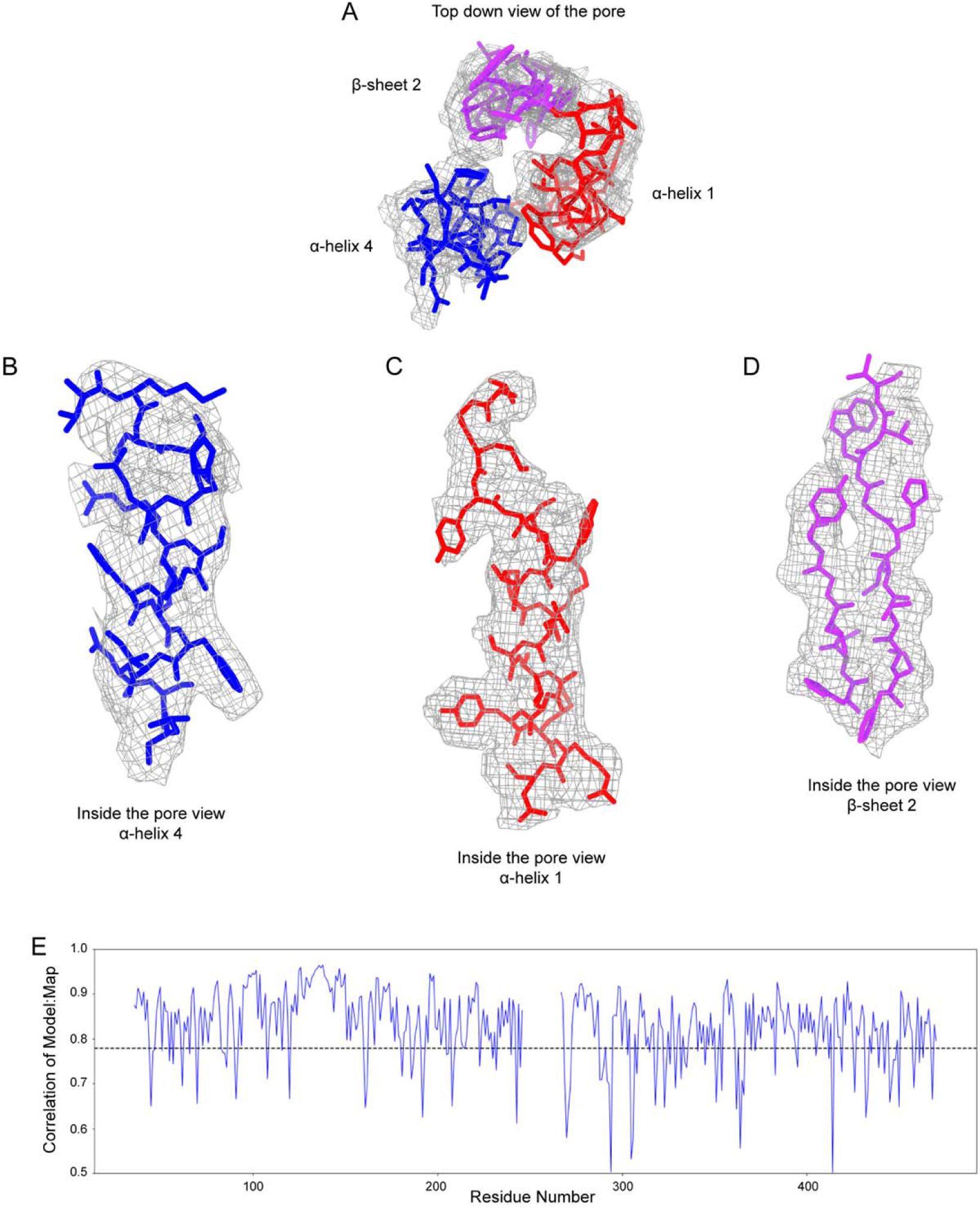
A) A view of the pore from the top with the three component secondary structures of the pore colored. B) The fit of α-helix 4 residues (blue) into the cryoEM density (mesh). C) The fit of α-helix 1 residues (red) into the cryoEM density (mesh). D) The fit of the first two strands of β-sheet 2 residues (purple) into the cryoEM density (mesh). E) The correlation between the cryoEM density and each residue of the model built into the cryoEM map. The average cross correlation of 0.77 is represented by a dotted line.

**Supplemental Figure 5:**
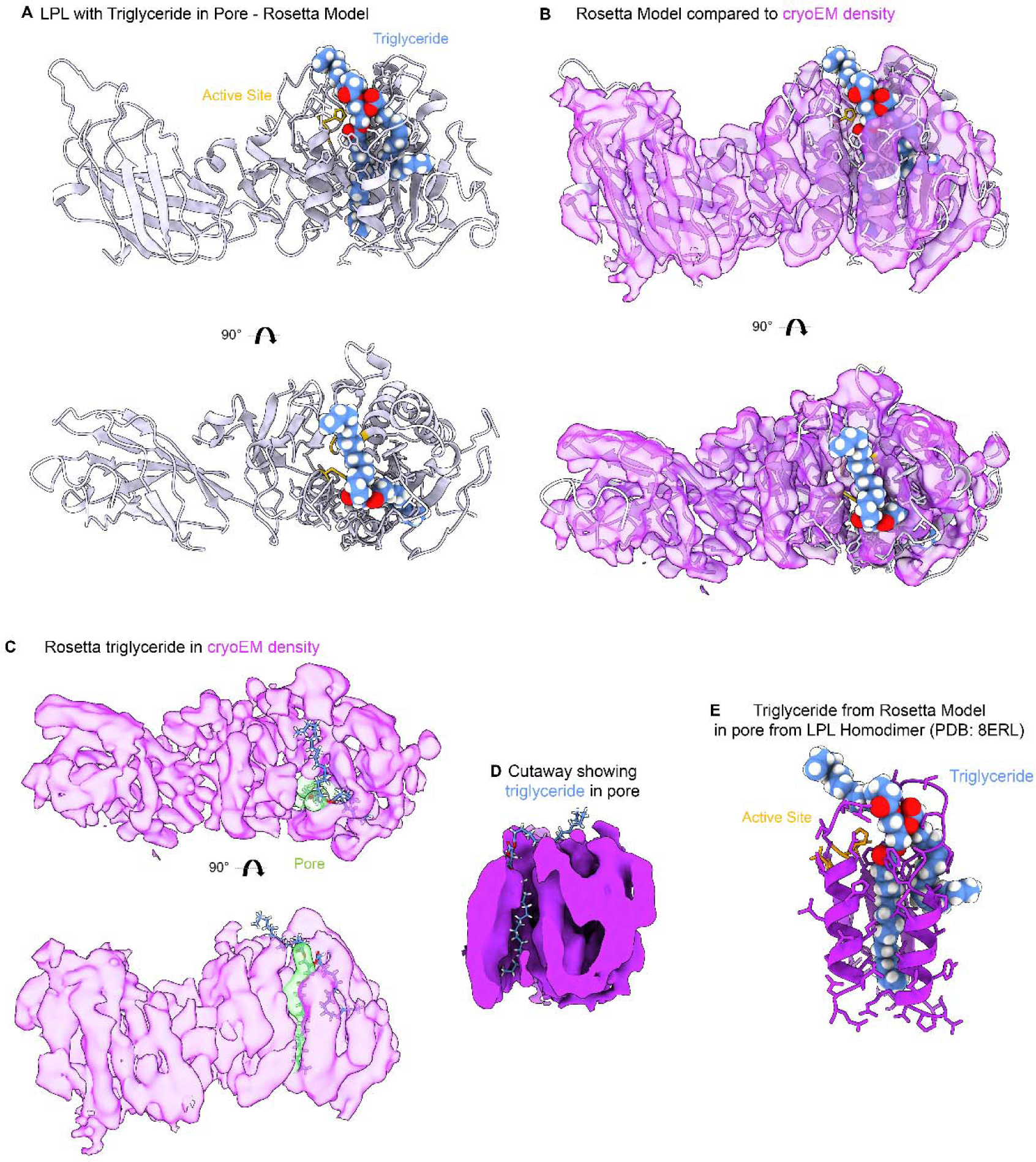
A) The results of PyRosetta modeling for LPL (white) with a triglyceride (light blue) ligand. The sn-1 acyl chain is in the hydrophobic pore, the sn-2 acyl chain is in a pocket to the side of the active site, the sn-3 acyl chain is in the hydrophobic pocket revealed by opening of the lid peptide. This model is available in supplemental files. B) Comparison of the PyRosetta model to the cryoEM density (transparent purple) C) The triglyceride from the PyRosetta model overlaid with the cryoEM map and identified pore (transparent lime green). D) A slice of the LPL density containing the triglyceride model showing the position of the sn1 chain in the pore and the sn-3 acyl chain in the upper hydrophobic pocket. E) The triglyceride fit in the pore residues from the LPL homodimer.

**Supplemental Figure 6:**
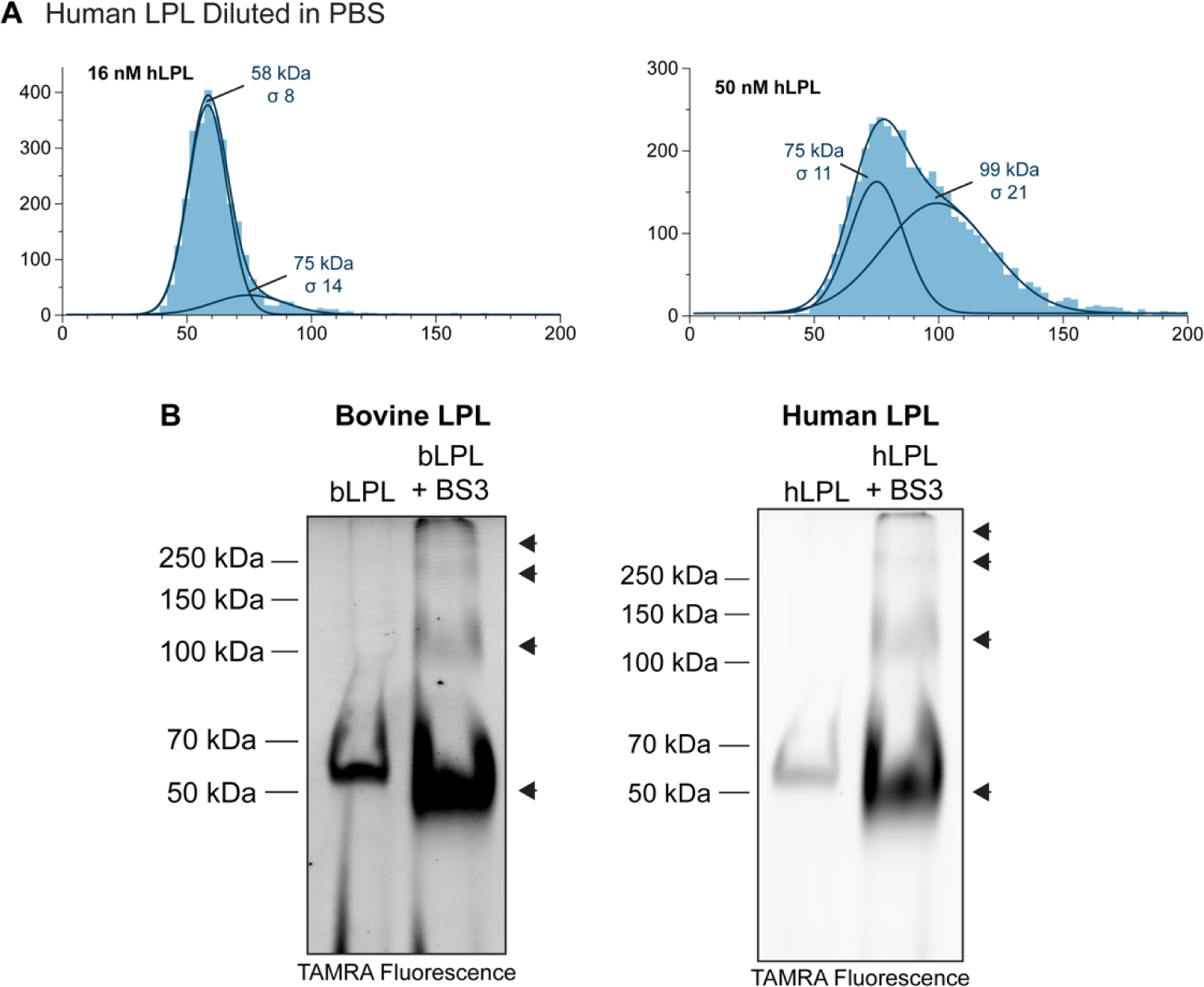
A) Human LPL (hLPL) (furin resistant mutant) was analyzed with mass photometry at 2 concentrations, 16 nM and 50 nM. Histogram of mass distributions were fit with multiple gaussians to determine an average molecular weight (kDa) and the width of the standard deviation of the fitted gaussian is given by sigma (σ). LPL monomer theoretical molecular weight = 50.5 kDa. At 16 nM hLPL the primary peak corresponds to the molecular weight of an LPL monomer (58 kDa). At 50 nM the predominate peak has shifted to 99 kDa, corresponding to a dimer of LPL. A lower molecular weight shoulder is seen (75 kDa), suggesting the shift to a completely dimerized population is incomplete. B) Crosslinking of both bovine LPL (bLPL) and hLPL with bis(sulfosuccinimidyl)suberate (BS3) analyzed by TAMRA-FP serine-hydrolyase probe fluorescence shows the appearance of at least 4 oligomeric species (arrows). Control uncrosslinked protein is in the first lane for comparison.

**Supplemental Figure 7:**
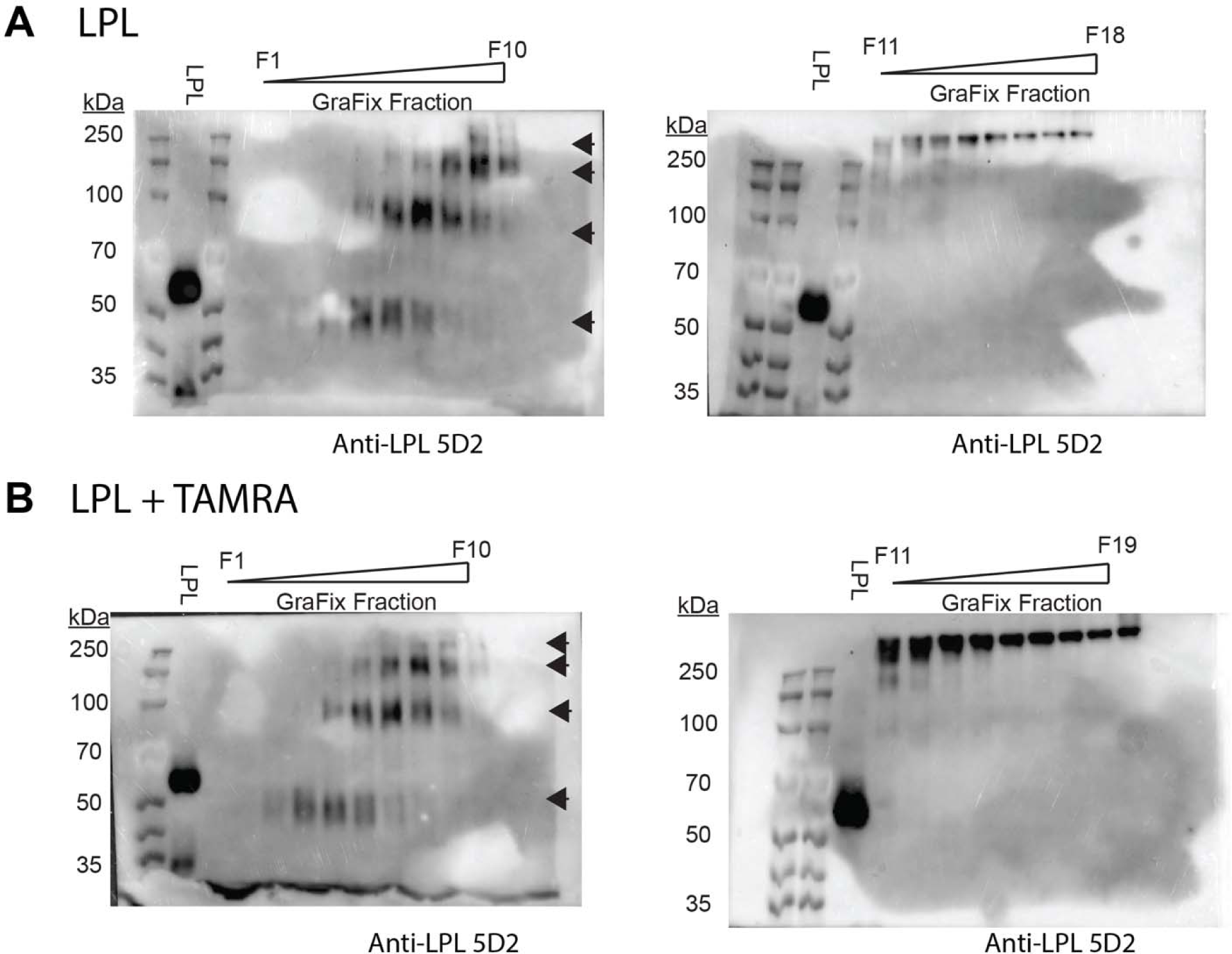
Addition of TAMRA-FP serine hydrolase probe (TAMRA) does not alter LPL oligomer distribution in GraFix. A) LPL crosslinked by GraFix has at least 4 distinct LPL oligomers (black arrows). B) LPL incubated with TAMRA and then run on GraFix shows the same oligomer formation. Western blots were developed with the LPL 5D2 antibody. Small differences in the oligomer distribution can be attributed to fractionating by hand. We subsequently used TAMRA fluorescence to identify LPL bands for downstream mass photometry due to its increased sensitivity and speed.

**Supplemental Figure 8:**
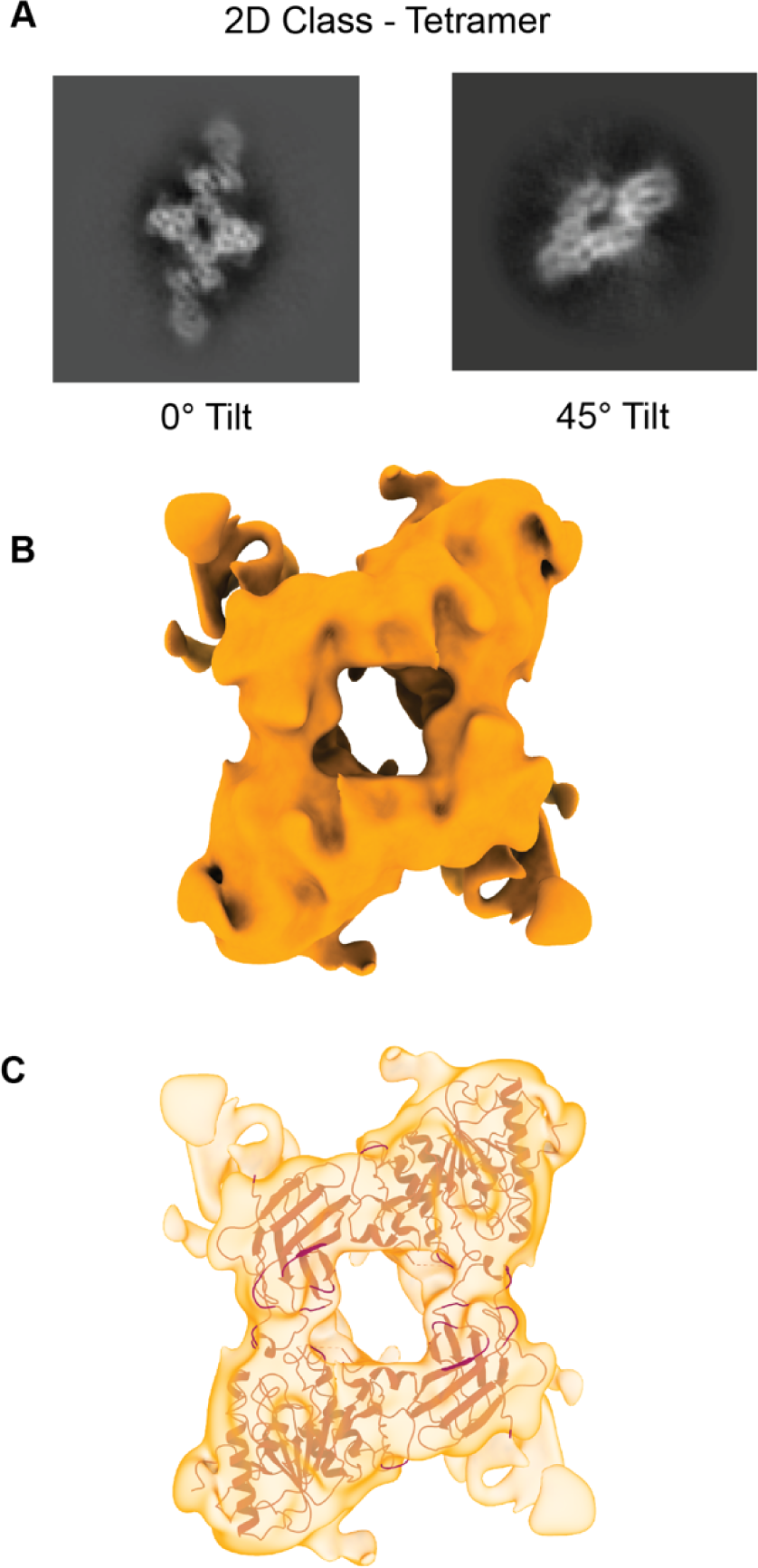
A) Some protein particles in our cryoEM data were observed adopting an apparently tetrameric oligomer of LPL – also in a preferred orientation – the center 2 interacting LPL subunits were the least mobile portion of the complex and therefore the easiest to resolve in 2D classification. We also observed the center of the tetramer in our tilted data. B) We generated a 3D model using cryosparc with the tetramer particles from selected 2D classes. C) The LPL dimer subunit of the LPL helix structure (PDB 6U7M) fits very well into the density suggesting that this tetramer interaction may be mediated by the 2 center LPL subunits adopting a structure like the inactive LPL dimer found in the helix.

**Supplemental Figure 9:**
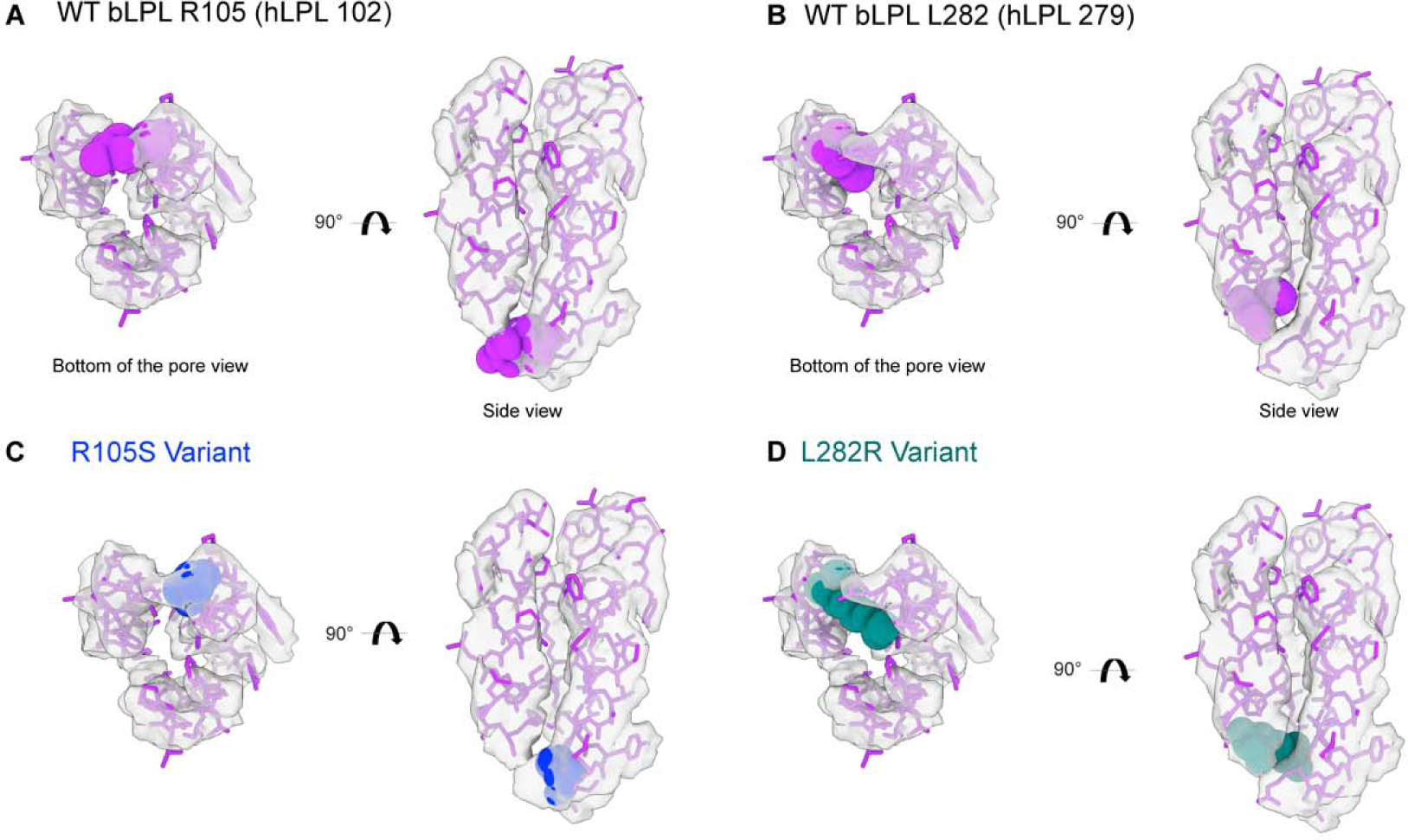
A) The residues (purple) and density (transparent gray) corresponding to the pore of LPL with R105 residue shown as a sphere representation (this corresponds to R102 in human LPL). B) The L282 residue shown as a sphere representation (this corresponds to L279 in human LPL). C) Modeling of the R105S variant (blue). D) Modeling of the L282R variant (teal).

**Supplemental Table 1:**
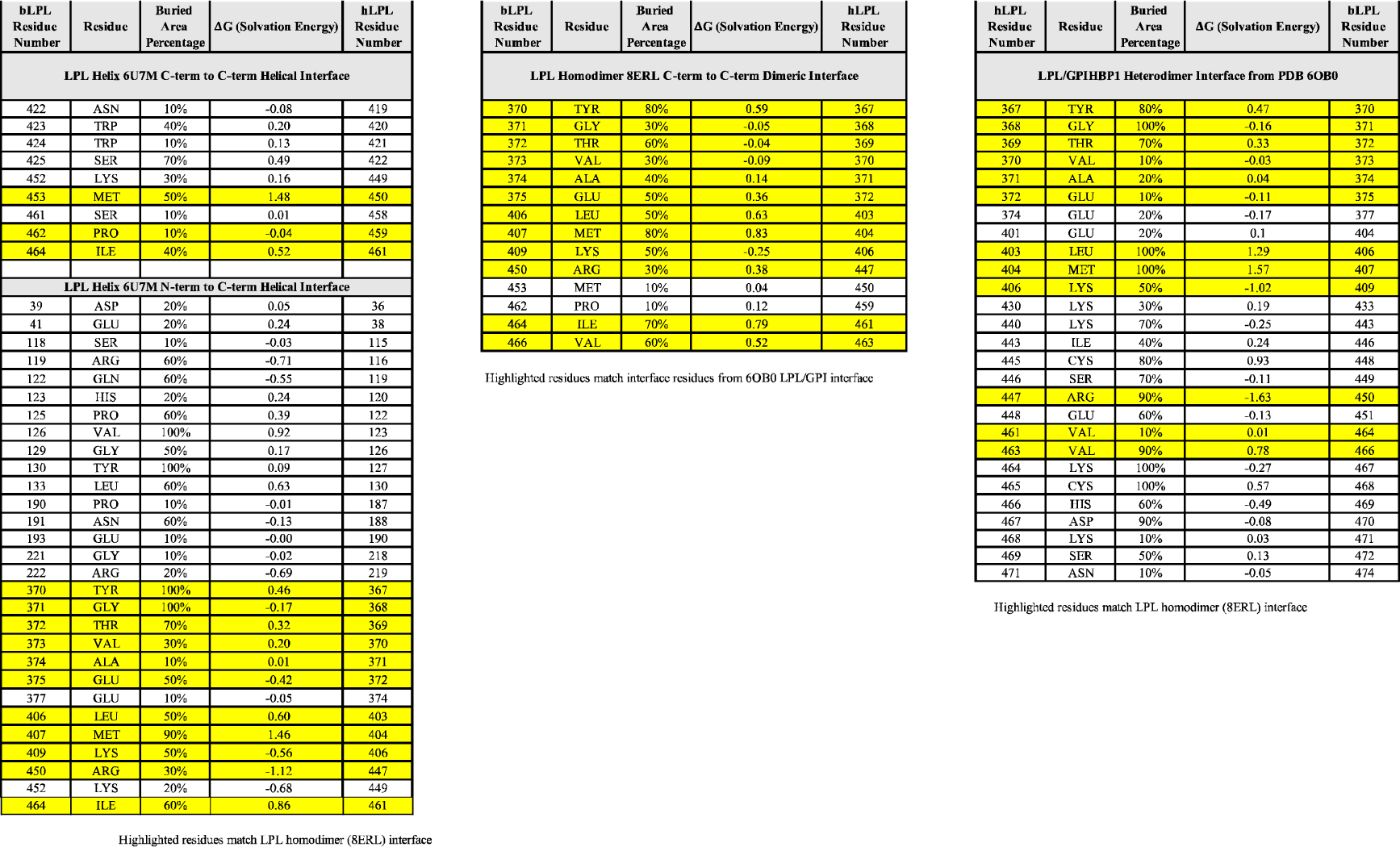
PDBePISA analysis results showing residues involved in interaction interfaces for the LPL helix (C-terminal to C-terminal and C-terminal to N-terminal), LPL homodimer (C-terminal to C-terminal), LPL/GPIHBP1 heterodimer (C-terminal to GPIHBP1). Highlighted residues indicate the overlap with the interface listed at the bottom of the table. Residue numbering for both human and bovine LPL is included to facilitate comparison.

## Supplemental Movie 1

A movie of a single LPL subunit morphing back and forth from the structure of LPL in the LPL/GPIHBP1 crystal structure (PDB 6OB0) to an LPL subunit from the LPL homodimer (PDB 8ERL). The opening of the active site pore in the LPL homodimer is the final state of the movie. This morph is not meant to represent the physiological movement that might occur during opening of the LPL pore, but meant to demonstrate the difference between these two structures of LPL solved by different methods, one with an active site inhibitor (6OB0) and one with an open pore (8ERL).

## Supplemental Movie 2

A movie of a single LPL subunit morphing back and forth from the structure of LPL from the LPL helical oligomer (PDB 6U7M) to an LPL subunit from the LPL homodimer (PDB 8ERL). The opening of the active site pore in the LPL homodimer is the final state of the movie. This morph is meant to demonstrate that the movement observed in supplemental movie 1 is similar to what we see when comparing two structures – one inactive (6U7M) and one active (8ERL) - that were solved by cryoEM.

## Supplemental Movie 3

A movie of a single LPL subunit morphing back and forth from the structure of LPL forth from the structure of LPL in the LPL/GPIHBP1 crystal structure (PDB 6OB0) to from the LPL helical oligomer (PDB 6U7M). This morph is meant to demonstrate that the movement observed in supplemental movies 1 is not necessarily a result of the technique or resolution, but the shifts observed in the structure going from an inactive to active state. This morph compares an inhibited structure (6OB0) to an inactive oligomer (6U7M)

## Supplemental Files

Files of the pdb files for the starting (labeled ‘start’) and ending (labeled ‘final’) models for fitting a palmitate (PLM), oleate (OLA), linoleate (EIC), and triglyceride (TGL) into LPL’s hydrophobic pore with PyRosetta. Excel files with the histograms of the mass photometry data from figure 5 and fitted gaussian curves for each.

